# Automated model discovery for textile structures: The unique mechanical signature of warp knitted fabrics

**DOI:** 10.1101/2024.07.26.605392

**Authors:** Jeremy A. McCulloch, Ellen Kuhl

**Affiliations:** Department of Mechanical Engineering, Stanford University, Stanford, California, United States

**Keywords:** textile structures, knitted fabrics, biaxial testing, constitutive modeling, anisotropy, constitutive neural networks, machine learning

## Abstract

Textile fabrics have unique mechanical properties, which make them ideal candidates for many engineering and medical applications: They are initially flexible, nonlinearly stiffening, and ultra-anisotropic. Various studies have characterized the response of textile structures to mechanical loading; yet, our understanding of their exceptional properties and functions remains incomplete. Here we integrate biaxial testing and constitutive neural networks to automatically discover the best model and parameters to characterize warp knitted polypropylene fabrics. We use experiments from different mounting orientations, and discover interpretable anisotropic models that perform well during both training and testing. Our study shows that constitutive models for warp knitted fabrics are highly sensitive to an accurate representation of the textile microstructure, and that models with three microstructural directions outperform classical orthotropic models with only two in-plane directions. Strikingly, out of 2^14^ =16,384 possible combinations of terms, we consistently discover models with two exponential linear fourth invariant terms that inherently capture the initial flexibility of the virgin mesh and the pronounced nonlinear stiffening as the loops of the mesh tighten. We anticipate that the tools we have developed and prototyped here will generalize naturally to other textile fabrics–woven or knitted, weft knit or warp knit, polymeric or metallic–and, ultimately, will enable the robust discovery of anisotropic constitutive models for a wide variety of textile structures. Beyond discovering constitutive models, we envision to exploit automated model discovery as a novel strategy for the generative material design of wearable devices, stretchable electronics, and smart fabrics, as programmable textile metamaterials with tunable properties and functions. Our source code, data, and examples are available at https://github.com/LivingMatterLab/CANN.

## 1. Motivation

Synthetic meshes have unique mechanical properties [3], which make them ideal candidates for many engineering and medical applications [19]: Under small deformations they display a remarkable *initial flexibility*, under large deformations, they exhibit a pronounced *nonlinear sti*ff*ening*, and under varying loading scenarios, they reveal an *ultra-anisotropic response* [18]. Their compliant nature allows them to effortlessly undergo large deformations while maintaining structural integrity [9]. Their inherent porosity encourages fluid flow and nutrient exchange, crucial for many biological or technical functions [41]. Their anisotropic nature enables them to adapt and function effectively in complex dynamic environments [38]. Collectively, these unique features make meshes highly suitable for a range of biomedical applications, including surgical repair, drug delivery, wound healing, implant design, and tissue engineering [37]. Beyond biomedicine, meshes are becoming increasingly popular in other technical applications including filtration systems, protective gear, flexible electronics, and structural reinforcement [29].

### The microstructure of the mesh is critical to its function

The mechanical properties of the mesh directly affect how well it integrates within its environment, distributes stresses, and provides structural support [7]. In fabric mechanics, we distinguish two fundamentally different types of meshes: woven and knitted [18]. The microstructure of *woven* fabrics consists of warp threads running lengthwise and shute or weft threads running widthwise. The shute threads are woven over and under the warp threads interlacing them at right angles to create a tight and stable grid-like pattern with two orthogonal microstructural directions, the warp direction ***w*** and the shute direction ***s***. Along these directions, woven fabrics are rather stiff; off-set at 45 degrees, they are fairly compliant, meaning they have a high tensile stiffness, but a low shear stiffness [44]. The microstructure of *knitted* fabrics consists of interlocking loops of yarn, creating a flexible and stretchable textile structure that can easily adapt its shape and undergo large deformations while maintaining structural integrity [24]. We distinguish two families of knitted structures: weft-knitted and warp knitted [21]. The microstructure of *weft knitted* fabrics consists of horizontal rows of interlocking loops. It is made by looping a single yarn back and forth horizontally across the fabric width, creating interconnected loops, row by row, with two orthogonal microstructural directions, the production or wale direction ***w*** and the yarn direction ***s***. Regardless of the direction of loading, weft knitted fabrics tend to be extremely extensible and dimensionally unstable [55]. The microstructure of *warp knitted* fabrics consists of vertical columns of interlocking loops. It is made by looping multiple parallel yarns vertically along the fabric length, zigzagging between two neighboring columns, creating interconnected loops, column by column, with the production or warp direction ***w*** and one or multiple yarn directions ***s***_*I*_ and ***s***_*II*_. In the warp direction, warp-knitted fabrics are fairly stiff; in the yarn directions, the stiffness is tunable by the interlocking pattern [50]. Of the three different microstructures in Figure 1, warp knitted structures are the most tunable: The design space of a single loop spans 3 × 2 × 3 = 18 possibile configurations, three for the incoming yarn, two for the loop itself–open or closed–, and three for the outgoing yarn [21]. For warp knitted structures made with two guide bars, this already results in 18 ×18 = 324 possible combinations, and modern knitting machines have four or more guide bars. This opens tremendous opportunities to design warp knitted fabrics with mechanically-guided microstructures and custom-designed properties [4].

**Figure 1:**
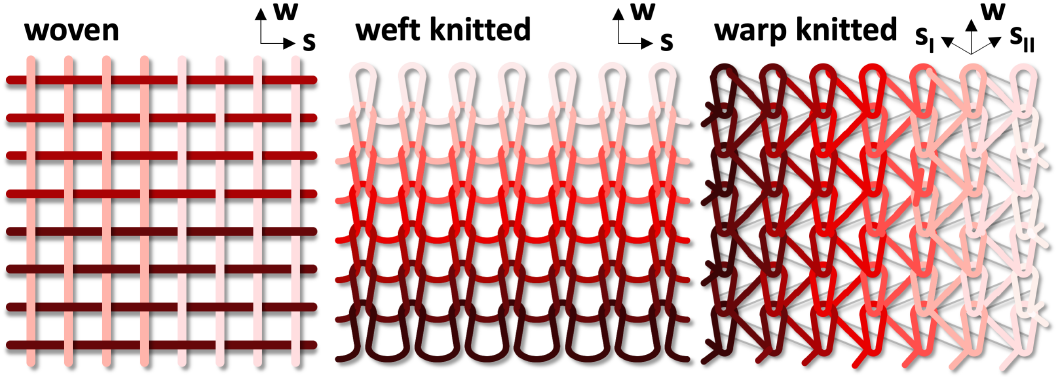
Microstructure of woven, weft and warp knitted fabrics. Woven fabrics consist of a series of parallel warp strands alternately passed over and under by a set of parallel shute strands creating a microstructure with two orthogonal directions, the warp direction ***w*** and the shute direction ***s***. Knitted fabrics consists of continuous filaments that are looped around one another. Weft knitted fabrics consists of horizontal rows of interlocking loops creating a microstructure with two orthogonal directions, the production or wale direction ***w*** and the yarn direction ***s***. Warp knitted fabrics consists of vertical columns of interlocking loops creating a microstructure with a warp direction ***w*** and one or more yarn directions ***s***_*I*_ and ***s***_*II*_.

### Synthetic meshes play an integral role in surgical repair

Initially introduced for hernia repair in the 1950s, the primary goal of synthetic meshes is to reinforce weakened or damaged structures and provide durable structural integrity. Hernia repair is a common surgery, with more than 20 million procedures each year [54]. In spite of this high prevalence, complications are relatively common, with 11 percent of patients reporting postoperative abdominal pain [42]. Hernia repairs use various methods to close the wound in the abdominal wall, but the most common surgical intervention is to suture a synthetic mesh over the wound [53]. In addition to the surface properties of the mesh at the mesh tissue interface that determine the durability of the reconstruction [23], it is essential that the surgeon selects a mesh with appropriate size, suture factors, and biomechanical properties [40]. A mismatch between the mechanics of the mesh and the surrounding tissue can increase the risk of postoperative failure [7]. To assess the mechanical biocompatibility of a synthetic mesh, we must understand its mechanical properties [13]. Thus, there is a critical need to develop constitutive models for synthetic meshes to better understand how they interact with the surrounding tissue [31]. Here, as a first step, we limit this understanding to the hyperelastic regime, and do not consider dynamic [20] or frictional [39] effects. Within this regime, previous studies have performed biaxial testing of various mesh patches [9]; however, these studies only extracted a few phenomenological parameters and did not combine data from multiple experiments into a unified mechanistic constitutive model. The benefit of obtaining a fully three-dimensional hyperelastic constitutive model from experimental data is that such a model would describe the stress state of the mesh induced by *any* possible deformation, including complex states that we do not test experimentally, but may well occur in vivo [51]. However, here we will *not* follow the traditional approach, propose yet another phenomenological model, and fit its parameters to data. Instead, *our objective is to discover a microstructurally motivated hyperelastic constitutive model for warp-knitted fabrics and establish a robust set of experimental and computational techniques to autonomously discover the best constitutive models and parameters for a variety of woven, weft and warp knitted textile structures and other synthetic microstructural materials*.

### Constitutive modeling requires deep expert knowledge

A constitutive model aims to predict the internal stress state of a material as a function of its strain. In particular, in developing elastic constitutive models, we assume that the Piola stress tensor is a function of the deformation gradient tensor [17]. There are various approaches to infer this relationship between the deformation gradient and the Piola stress from mechanical testing data. Classical approaches attempt to fit a few unknown parameters of well-known phenomenological models such as the neo Hooke model [60], the Lanir model [22], or the Demiray model [8]. The challenge with this approach is that it requires a profound domain expertise to select an appropriate model, and we may need to repeat model selection and parameter identification several times to identify a model that fits the data well. Constitutive neural networks use physics-informed insights about constitutive models as well as powerful deep learning methods to simultaneously discover the function form of the constitutive model and learn its appropriate parameters [26]. While the first family of constitutive neural networks only discovered isotropic models for materials such as rubber [26], brain [27] or plant-based meat [47], more recent networks can now discover transversely isotropic models for skin [28] or arteries [45]. Here we explore how to expand constitutive neural networks to discover more complex anistropic constitutive models from biaxial extension data.

### Constitutive neural networks can autonomously discover the best model and parameters

A constitutive neural network is a machine learning model that takes the deformation gradient as an input and outputs the Piola stress for a specific material [25]. Constitutitve artificial neural networks take advantage of physical laws and assumptions about the material to constrain the input-output relationship that the network discovers, similar to physics-informed neural networks [49]. However, constitutive neural networks differ from physics-informed neural networks, which alter the loss function to satisfy physical laws. Instead, constitutive neural networks modify the architecture of the underlying machine learning model such that, regardless of the model parameters, the discovered model exactly satisfies the relevant physical laws [26]. The particular assumptions that motivate the design of hyperelastic constitutive neural networks are material objectivity, thermodynamic consistency, incompressibility, and material symmetry. Prior work has enforced various forms of symmetry including isotropy [26], transverse isotropy [28], and orthotropy [30]; here we assume that warp-knitted meshes have either two or three characteristic microstructural orientations in the plane of the mesh.

Unlike most previous work that trains anisotropic constitutive neural networks exclusively by using homogeneous tests [58], we explore how to use biaxial testing data when neither of the loading axes are aligned with the symmetry planes of the material [52]. We acknowledge that this violates the condition of homogeneity; yet, it allows us to train our network with a broader class of deformations, and discover models that are more robust when predicting stresses in response to deformation states that the network has not previously seen during training [59]

Finally, while constitutive neural networks are able to discover constitutive models by selecting from a large set of possible strain energy functions, the most useful constitutive models have only a small number of parameters and are therefore interpretable [5]. Various regularization techniques can help reduce the number of non-zero parameters without significantly sacrificing model accuracy [14]. Prior work has shown that, when computationally tractable, *L*_0_ regularization is the best method for identifying the optimal *n*-term model for a fixed, small *n* [32]. When *n* is too large for *L*_0_ optimization, *L*_*p*_ regularization with 0 < *p*≤1 can effectively reduce the number of non-zero terms [46], without significantly decreasing model accuracy [12]. Here we will apply both approaches, compare the discovered models, and make recommendations which regularization to choose.

## 2. Methods

### 2.1. Experimental methods

We prepared samples from a 0.5 mm thick warp knitted surgical mesh of extruded polypropylene designed for surgical repair (PROLENE^OR^ Ethicon, Inc., Somerville, NJ), to test under biaxial loading in a CellScale BioTester 5000. We define the *warp direction* of the mesh as the direction along which the loops are aligned, which is also the direction in which the mesh is stiffest. We then define the *shute directions* in the plane of the mesh, either as one direction orthogonal to the warp direction inclined by 90 degrees, or as two directions symmetrically inclined to the warp direction by 60 degrees. We cut the mesh into square sections two different orientations: In the first orientation, which we denote as the *0*/*90 orientation* or *0*/+*60*/*-60 orientation*, the sample is aligned with the warp direction; in the second orientation, which we denote as the *-45*/+*45 orientation* or *-45*/+*15*/+*75 orientation*, the sides of the sample are 45 degrees offset from the warp direction. Figure 2 illustrates the samples of each orientation, mounted into the testing device. In the 0/90 or 0/+60/-60 orientation, in the left column, we place the tines of the biaxial testing device in the loops of the mesh. To prevent the mesh from unravelling, we avoid placing the tines in the row of loops that is closest to the edge of the sample. In the -45/+45 or -45/+15/+75 orientation, in the right column, we place the tines in the loops of the mesh where feasible, but due to the geometry of the mesh not all tines fit exactly into a loop. To reduce the likelihood of unraveling, we use a larger mesh area in this orientation.

**Figure 2:**
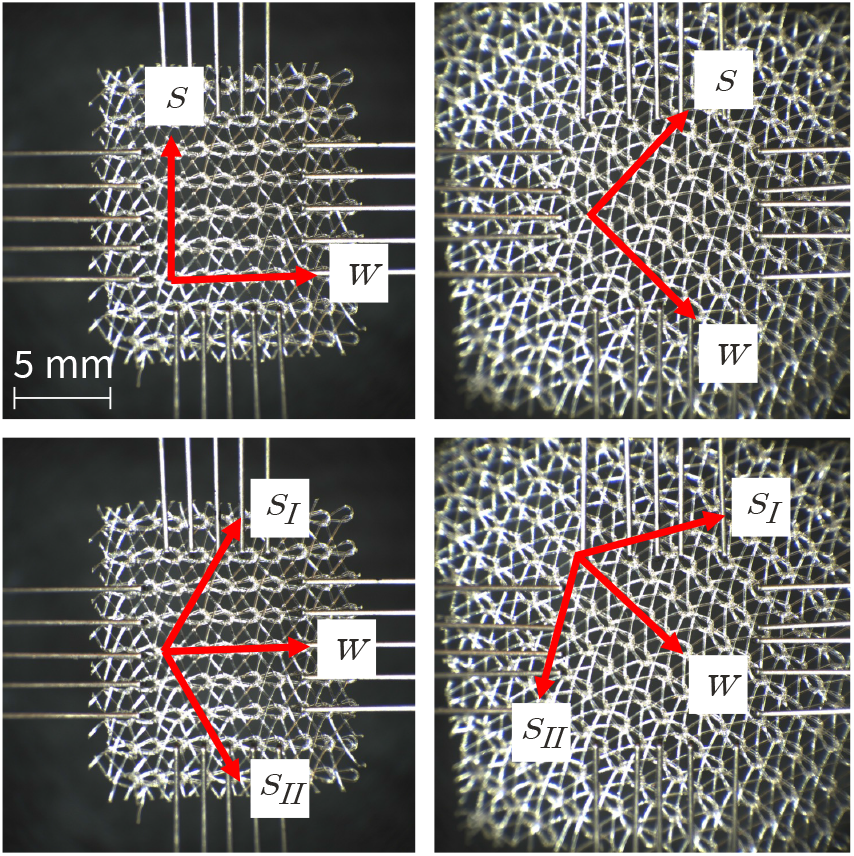
Biaxial testing of the mesh in two orientations. The top row shows the the two-fiber model with one warp and one shute direction ***w*** and ***s***, and the bottom row shows the three-fiber model with one warp and two shute directions ***w*** and ***s***_*I*_ and ***s***_*II*_. The left column illustrates the mesh in the 0/90 or 0/+60/-60 orientations, and the right column illustrates the mesh in the -45/+45 or -45/+15/+75 orientations.

We test a total of ten samples, five for each orientation [10]. Once we mount each sample into the device, we apply a 30 mN preload along both axes to establish a consistent reference configuration [61]. After preloading, we record the spacing between the tines as the gauge length, and perform five consecutive tests with 30 seconds of rest between each test [33]. Each test consists of linearly increasing the applied stretch for 100 seconds, followed by linearly decreasing the applied stretch for 100 seconds until the sample returns to its unstretched state. Table 1 summarizes the maximum stretches in each direction for each of the five tests. We perform all five tests on each of the ten samples. However, to minimize the effect of preload, we randomize the order of the tests for each sample.

**Table 1:** Definition of five biaxial test settings. Maximum stretches *λ*_1_ and *λ*_2_ for the five different tests of each sample. We increase the stretch linearly for 100 seconds until reaching the maximum stretch for each test, then decrease it linearly for 100 seconds to the undeformed configuration, and hold the sample in the undeformed configuration for 30 seconds between consecutive tests. To minimize the effect of preload, we randomize the order of the tests for each sample.

| experiment | max stretch $\lambda_1$ | max stretch $\lambda_2$ |
| --- | --- | --- |
| strip-x | 1.10 | 1.00 |
| off-x | 1.10 | 1.05 |
| equibiax | 1.10 | 1.10 |
| off-y | 1.05 | 1.10 |
| strip-y | 1.00 | 1.10 |

### 2.2. Stress and strain analysis

We process the data from the CellScale BioTester 5000 to obtain the average Piola stress for given stretch values *λ*_1_ and *λ*_2_. First, we convert the force and displacement measurements to stresses and stretches. To do so, we measured the sample thickness *t* with calipers and set the gauge lengths *L*_1_ and *L*_2_ to the initial spacing between the tines in the 1- and 2-directions. Then, we computed the stretches *λ*_1_ and *λ*_2_ and the Piola stresses *P*_11_ and *P*_22_,

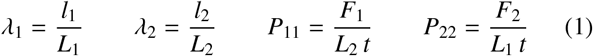

where *l*_1_ and *l*_2_ are the measured gauge lengths and *F*_1_ and *F*_2_ are the measured forces in the 1- and 2-directions. The result are five loading and five unloading curves for each of the ten samples. We resample and average all curves at equidistant stretch intervals to obtain an averaged stress pair {*P*_11_, *P*_22_} for each stretch pair {*λ*_1_, *λ*_2_}. Finally, for the samples mounted in the -45/+45 orientation, we use symmetry with respect to the diagonale and average the ten recorded stress-stretch curves to five distinct curves.

### 2.3. Kinematics

We characterize the deformation through the mapping ***x*** = ***ϕ***(***X***) that maps a point ***X*** in the reference configuration to a point ***x*** the deformed configuration. We then describe the local deformation using the deformation gradient,

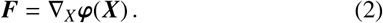

Multiplying ***F*** with its transpose ***F***^t^ introduces the symmetric right Cauchy Green deformation tensor ***C***,

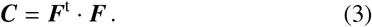

To characterize the deformation of an orthotropic material, we introduce the three isotropic invariants, *I*_1_, *I*_2_, *I*_3_, and a set of anisotropic invariants, *I*_4*i*_, *I*_8*i j*_ [56, 35],

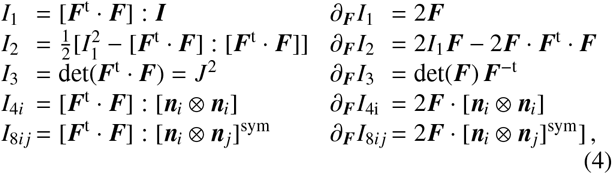

where ***I*** is the identity tensor, ***n***_*i, j*_ with *i, j* = 1, …, *n*_dir_ are the unit warp and shute directions in the undeformed reference configuration, and *n*_dir_ is the number of microstructural directions of our model. Since we are unable to measure the deformation in the thickness direction of the mesh, we assume that the mesh is perfectly incompressible, *I*_3_ = 1.

In our biaxial extension tests, we stretch the sample in two orthogonal directions, *λ*_1_ ≥ 1 and *λ*_2_ ≥ 1. The incompressibility condition, 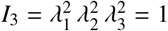, defines the stretch in the thickness direction as *λ*_3_ = (*λ*_1_ *λ*_2_)^−1^ ≤1. We assume that the deformation remains homogeneous and shear free, and the deformation gradient ***F*** remains diagonal,

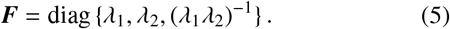

Next, we specify the invariants (4) in terms of the stretches *λ*_1_ and *λ*_2_ for two the microstructural architectures in Figures 1 and 2, the two-fiber model and the three-fiber model.

#### 2.3.1. Two-fiber model

For the two-fiber model, we assume that we can represent the mesh through two orthogonal fiber families, one in the warp direction ***w***, and one in the shute direction ***s***, inclined against the warp direction by π/2 = 90^°^,

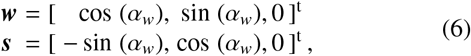

where α_*w*_ is the warp angle, the angle of the warp direction against the 1-direction. We now specify the invariants (4) in terms of the biaxial stretches *λ*_1_ and *λ*_2_ and the warp angle α_*w*_,

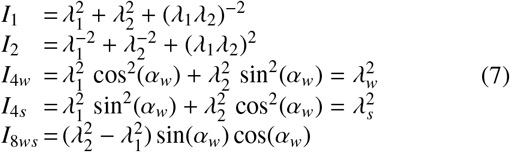

and take their derivatives,

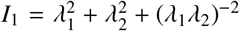

Our two-fiber model has five distinct invariants, *I*_1_, *I*_2_, *I*_4*w*_, *I*_4*s*_, *I*_8*ws*_, where *I*_4*w*_ and *I*_4*s*_ take the interpretation of the squared stretches in the warp and shute directions *λ*_*w*_ and *λ*_*s*_, and *I*_8*ws*_ characterizes the interaction between the warp and shute directions. For the special case of the 0/90 orientation, the anisotropic invariants are

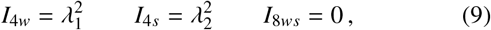

and we already see that this setup is incapable of probing the interaction between the warp and shute directions since *I*_8*ws*_ = 0. For the special case of the +45/-45 orientation, the anisotropic invariants are

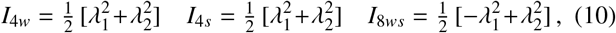

and we already see that this setup is incapable of distinguishing between the warp and shute directions since *I*_4*w*_ = *I*_4*s*_.

#### 2.3.2. Three-fiber model

The potential shortcomings of the two-fiber model motivate a more advanced three-fiber model with three fiber families, one in the warp direction ***w***, and two in the shute directions ***s***_*I*_ and ***s***_*II*_, symmetrically inclined against the warp direction by by +π/3 = +60^°^ and −π/3 = −60^°^,

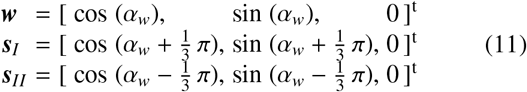

where α_*w*_ denotes the warp angle against the 1-direction. We now specify the invariants (4) in terms of the biaxial stretches *λ*_1_ and *λ*_2_ and the warp angle α_*w*_,

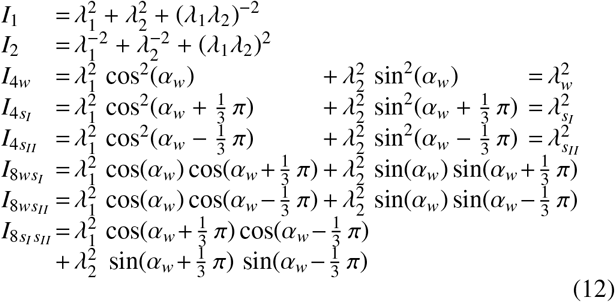

Our three-fiber model has eight distinct invariants, 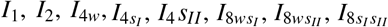, where 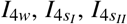 take the interpretation of the squared stretches in the warp and shute directions 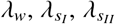 and 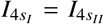 characterizes the interaction between the warp and shute directions. For the special case of the 0/+60/-60 orientation, the anisotropic invariants are

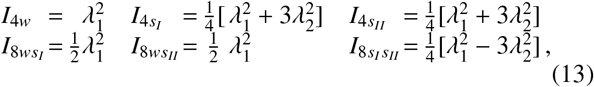

and we notice the microstructural symmetry of the two shute directions as 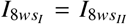 and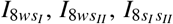. For the special case of the -45/+15/+75 orientation, the anisotropic invariants are

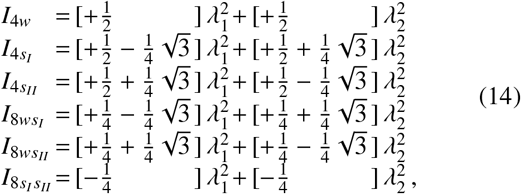

and we immediately notice that the three-fiber model invariants are much richer than the two-fiber model invariants.

### 2.4. Constitutive modeling

A constitutive model is a material-specific function that estimates the stress in a material given its deformation history. In the case of a hyperelastic constitutive model, the stress, in our case the Piola stress ***P***, only depends on the current deformation state, in our case the deformation gradient ***F***, such that ***P*** = ***P***(***F***). To satisfy thermodynamic consistency, we can express the stress as a function of the strain energy density *ψ* as ***P*** = *∂ψ*(***F***)/*∂****F***, which we reformulate in terms of our set of invariants *ψ*(*I*_1_, *I*_2_, *I*_3_, *I*_4*i*_, *I*_8*i j*_),

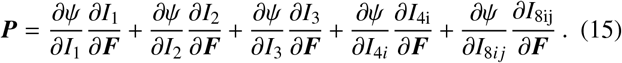

where *i, j* = 1, …, *n*_dir_ is the number of microstructural directions of our model. We explicitly enforce incompressibility by selecting the term in the third invariant as *ψ*(*I*_3_) = −*p* [*J*−1], such that *∂ψ*/*∂I*_3_ *∂I*_3_/*∂****F*** = −*p* ***F***^−t^. Here *p* acts as a Lagrange multiplier that we determine from the zero-thickness-stress condition.

### 2.5. Biaxial testing

In biaxial extension tests, we stretch the sample in two orthogonal directions, *λ*_1_ ≥ 1 and *λ*_2_ ≥ 1, and, by incompressibility, *λ*_3_ = (*λ*_1_ *λ*_2_)^−1^ ≤1. We assume that the deformation remains homogeneous and shear free, and that the resulting Piola stress ***P*** remains diagonal,

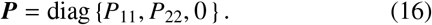

We use the isotropic first and second invariants 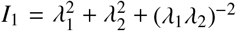 and 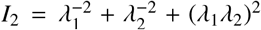 from equation (7) and their derivatives from equation (8) to determine the pressure *p* from the zero-thickness-stress condition in the third direction,

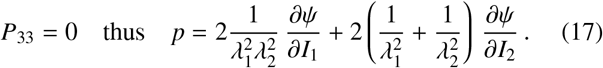

Equations (15) and (17) then provide an explicit analytical expression for the nominal stresses *P*_11_ and *P*_22_ in terms of the stretches *λ*_1_ and *λ*_2_ and the warp angle α_*w*_. We acknowledge that biaxial testing is limited to characterize the stress-stretch behavior within the plane, and does not allow us to make any assessment of the out-of-plane behavior of the fabric.

#### 2.5.1. Two-fiber model

For the two-fiber model, the free energy function is a function of five terms, *ψ* = *ψ*(*I*_1_, *I*_2_, *I*_4*w*_, *I*_4*s*_, *I*_8*ws*_), and the nominal stresses *P*_11_ and *P*_22_ have five terms, one for each invariant,

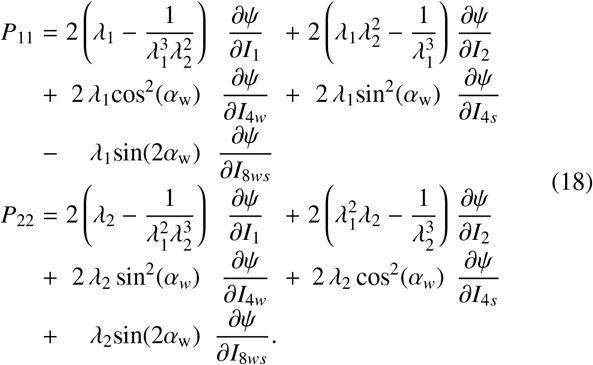

Here we make an important observation: Counterintuitively, the eighth-invariant term, *∂ψ*/*∂I*_8*ws*_, contributes negatively to the stresses in the 1-direction and positively to the stresses in the 2-direction. This contradicts our general intuition, and potentially violates the condition of material symmetry. We therefore drop the dependence on the eighth invariant and reduce our set of invariants to four, *ψ* = *ψ*(*I*_1_, *I*_2_, *I*_4*w*_, *I*_4*s*_). For the special case of the 0/90 orientation, with α_*w*_ = 0 π = 0^°^, the biaxial Piola stresses simplify to

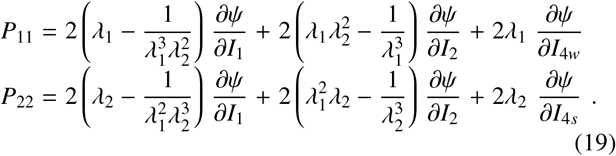

For the special case of the -45/+45 orientation, with α_*w*_ = −π/4 = −45^°^, the stresses are

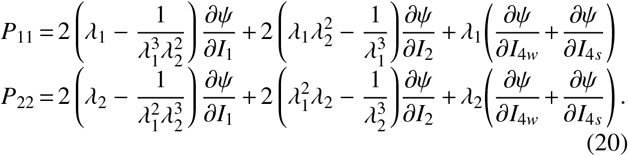

#### 2.5.2. Three-fiber model

For the three-fiber model, for similar reasons as above, we drop the dependence on the eighth invariants and reduce our set of invariants to five,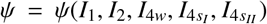. We now express the fourth invariants of the three-fiber model, 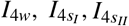, in terms of the invariants of the two-fiber model, *I*_4*w*_, *I*_4*s*_, *I*_8*ws*_, *I*_4(°)_ = *I*_4*w*_ cos^2^(α_°_) + *I*_*s*_ sin^2^(α_°_) + *I*_8*ws*_ sin(2α_°_). For our specific three-fiber model, with (°) = { *w, s*_*I*_, *s*_*II*_}, and 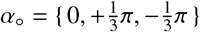, this allows us express the nominal stresses *P*_11_ and *P*_22_ using equation (18),

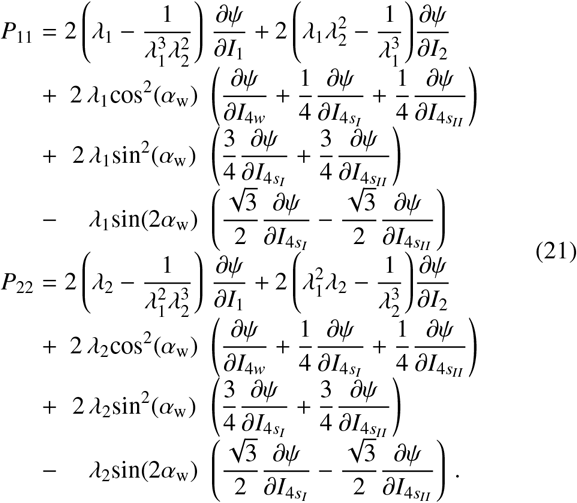

For the special case of the 0/+60/-60 orientation, with α_*w*_ = 0 π = 0^°^, the biaxial Piola stresses become

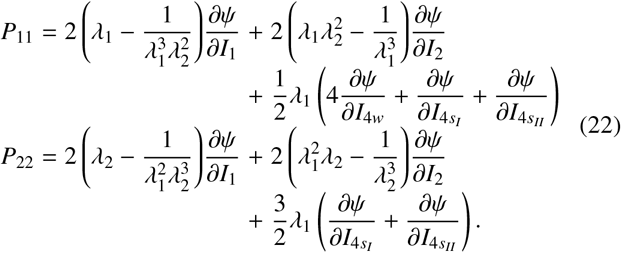

For the special case of the -45/+15/+75 orientation, with α_*w*_ = −π/4 = −45^°^, the biaxial Piola stresses are

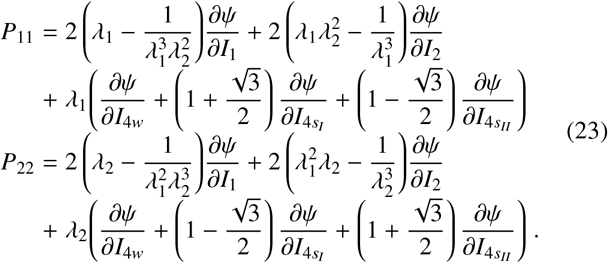

### 2.6. Constitutive neural networks

Motivated by the previous considerations, we design two constitutive neural networks to learn the free energy function *ψ*, a two-fiber network based on four invariants and a three-fiber network based on five invariants.

#### 2.6.1. Two-fiber network

Our first neural network approximates a strain energy function in terms of four invariants *I*_1_, *I*_2_, *I*_4*w*_, *I*_4*s*_ and has 28 parameters or network weights, 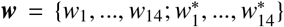 internal weights 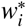 between its two hidden layers and 14 external weights *w*_*i*_ out of its final hidden layer. We assume that the individual contributions to the free energy are fully decoupled. For the isotropic terms *ψ*(*I*_1_) and *ψ*(*I*_2_), we adapt an isotropic constitutive neural network initially designed for rubber-like materials [26]. For the anisotropic terms *ψ*(*I*_4*w*_) and *ψ*(*I*_4*s*_), we adapt an anisotropic constitutive neural network initially designed for arteries [45], with the additional constraint that the linear and exponential linear terms in the warp direction, 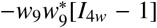 and 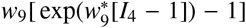, and in the shute direction, 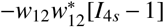 and 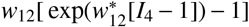, are not independent but share the same weights [63]. The free energy function for the two-fiber network takes the following explicit form,

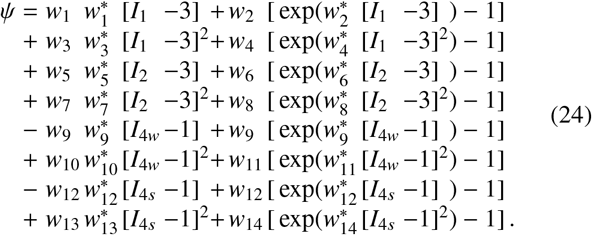

Figure 3 summarizes the architecture of the two-fiber family constitutive neural network.

**Figure 3:**
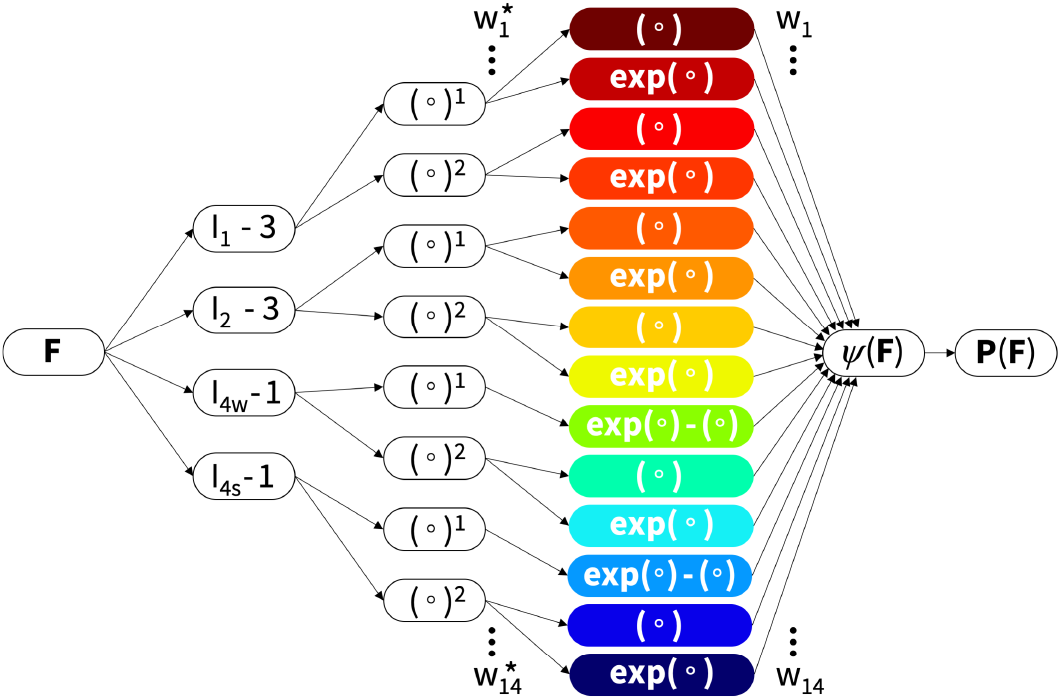
Two-fiber constitutive neural network. The network takes the two isotropic invariants *I*_1_ and *I*_2_ and the two anisotropic invariants *I*_4*w*_ and *I*_4*s*_ of the warp and shute directions ***w*** and ***s*** as input. The first layer generates powers (°) and (°)^2^ of the input and the second layer applies the identity (°) and exponential function (exp(°)) to these powers. The network learns the free energy function *ψ* as the weighted sum of the final layer, from which it derives the stress ***P***.

#### 2.6.2. Three-fiber architecture

Our second model architecture approximates a strain energy function in terms of five invariants *I*_1_, *I*_2_, *I*_4*w*_, *I*_4*sI*_, *I*_4*sII*_ and has 28 parameters or network weights, 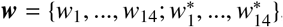, 14 internal weights 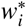 between its two hidden layers and 14 external weights *w*_*i*_ out of its final hidden layer. We use a similar neural network as before, and assume that the two shute directions ***s***_*I*_ and *s*_*II*_ have the same microstructure and share the same weights *w*_12_, *w*_13_, *w*_14_ and 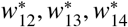. The free energy function for the three-fiber network takes the following explicit form,

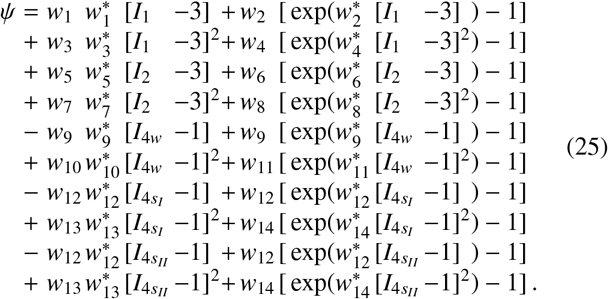

Figure 4 summarizes the architecture of the three-fiber family constitutive neural network.

**Figure 4:**
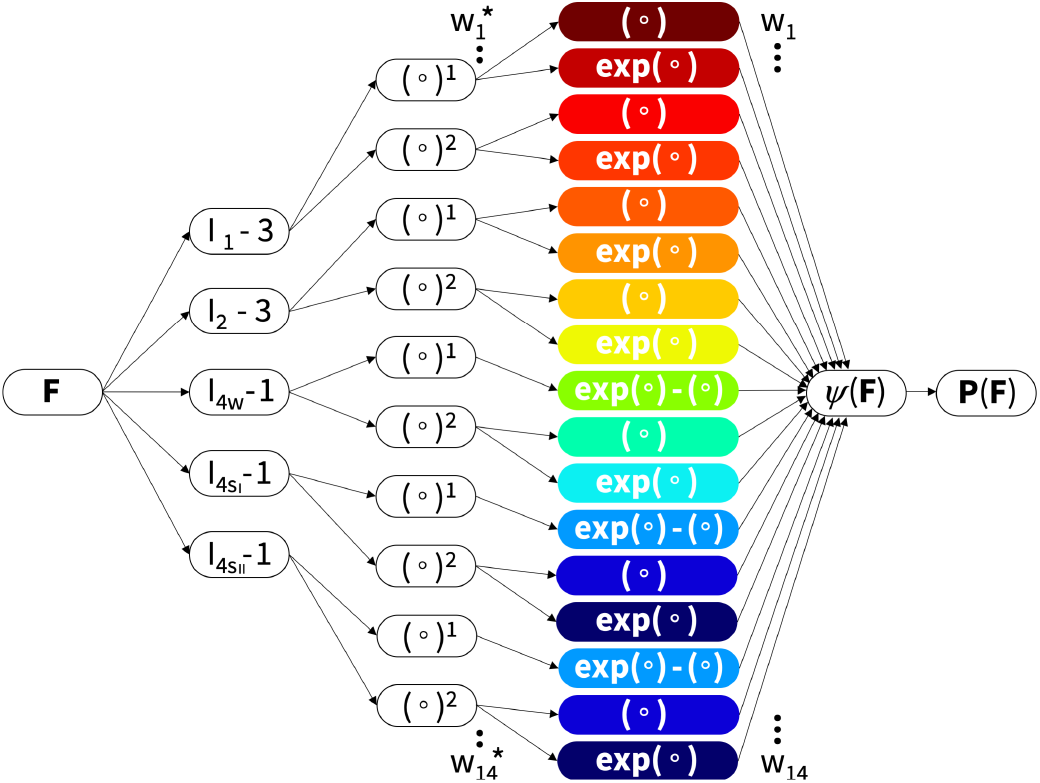
Three-fiber constitutive neural network. The network takes the two isotropic invariants *I*_1_ and *I*_2_ and the three anisotropic invariants *I*_4*w*_ and 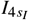 and 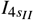 of the warp and shute directions ***w*** and ***s***_*I*_ and ***s***_*II*_ as input. The first layer generates powers (°) and (°)^2^ of the input and the second layer applies the identity (°) and exponential function (exp(°)) to these powers. The network learns the free energy function *ψ* as the weighted sum of the final layer, from which it derives the stress ***P***. We assume that the two shute directions ***s***_*I*_ and *s*_*II*_ have the same microstructure and share the same weights.

### 2.7. Model training

To discover models and parameters 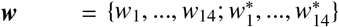 that best describe our synthetic mesh, we use the Adam optimizer to perform gradient descent on a weighted least squared error loss function *L* that penalizes the error between the discovered model ***P***(***F***_*i*_, ***w***) and the experimental data 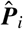 at *i* = 1, …, *n*_data_ discrete points, supplemented by *L*_*p*_ regularization,

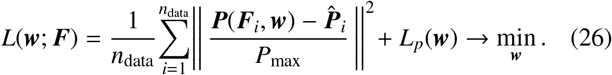

To account for all experiments equally, we weigh the error of each of the stress-stretch curve by the inverse of its maximum stress *P*_max_ [32]. For the *L*_0_ regularization, we supplement the loss function by an *α*-weighted regularization term, *L*_0_ = *α* || ***w*** ||_0_ with 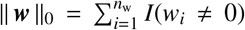, where *I*(°) is the indicator function that is one if the condition inside the parenthesis is true and zero otherwise. In practice, instead of solving the full discrete combinatorial problem and exploring all possible 2^14^ = 16, 384 combinations of terms, we only explore the weights and losses of the possible 14 one-term models and 91 two-term models by explicitly setting all other terms to zero. For the *L*_0.5_ regularization, we supplement the loss function by an *α*-weighted regularization, 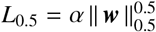 with 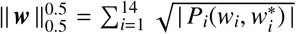, where *P*_*i*_ is the color-coded area of the stress contribution of the *i*-th term across all loading modes that we compute by summing the strain energies at the maximum displacement for each loading mode. The penalty parameter *α* sets the strength of the *L*_0.5_ regularization. We increase *α* progressively until the fit of the model to the data starts to become noticeably worse. To minimize the training time and ensure that the initial model creates predictions that are of the same order of magnitude as the data, we initialize all weights with a uniform distribution ***w*** =∼ Uniform([0, γ)), where we choose γ such that, in expectation, the total area under the measured stress-stretch curve equals the total area under the predicted stress-stretch curve.

## 3. Results

We perform all experiments as described in Section 2.1 and examine the data for each individual experiment. Figures 5 and 6 show the loading and unloading curves for the 0/90 orientation in the top two rows and for the -45/+45 orientation in the bottom row. The solid lines represent the means of *n* = 5 tests and the shaded areas represent the standard deviations. In all experiments, we observe significant hysteresis between the loading and unloading curves. The stress remains constant during the thirty-second holding between each experiment, from which we conclude that the difference in loading and unloading is a result of microstructural rearrangements of the mesh, rather than a viscous effect. For model discovery, for each experiment type, we extract the stretches and average stresses across the *n* = 5 tests during loading and unloading and summarize the stretch-stress data of all ten experiments in Table 2.

**Table 2:** Biaxial test data. Stress-stretch data for strip-x, off-x, equibiax, off-y, and strip-y tests in the 0/90 orientation, top, and in the -45/+45 orientation, bottom. Stress values are the means of *n* = 5 loading and unloading tests.

| 0/90 orientation<br>strip-w |  |  | 0/90 orientation<br>off-w |  |  | 0/90 orientation<br>equi-biax |  |  | 0/90 orientation<br>off-s |  |  | 0/90 orientation<br>strip-s |  |  |
| --- | --- | --- | --- | --- | --- | --- | --- | --- | --- | --- | --- | --- | --- | --- |
| $\lambda_w : \lambda_s = 1.10 : 1.00$ | | | $\lambda_w : \lambda_s = 1.10 : 1.05$ | | | $\lambda_w : \lambda_s = 1.10 : 1.10$ | | | $\lambda_w : \lambda_s = 1.05 : 1.10$ | | | $\lambda_w : \lambda_s = 1.00 : 1.10$ | | |
| $\lambda_w$<br>[-] | $\sigma_w$<br>[kPa] | $\sigma_s$<br>[kPa] | $\lambda_w$<br>[-] | $\sigma_w$<br>[kPa] | $\sigma_s$<br>[kPa] | $\lambda_w$<br>[-] | $\sigma_w$<br>[kPa] | $\sigma_s$<br>[kPa] | $\lambda_s$<br>[-] | $\sigma_w$<br>[kPa] | $\sigma_s$<br>[kPa] | $\lambda_s$<br>[-] | $\sigma_w$<br>[kPa] | $\sigma_s$<br>[kPa] |
| 1.000 | 0.00 | 0.00 | 1.000 | 0.00 | 0.00 | 1.000 | 0.00 | 0.00 | 1.000 | 0.00 | 0.00 | 1.000 | 0.00 | 0.00 |
| 1.006 | 4.49 | 1.59 | 1.006 | 4.88 | 6.88 | 1.006 | 5.68 | 10.01 | 1.006 | 2.82 | 8.59 | 1.006 | 1.76 | 10.05 |
| 1.013 | 7.12 | 2.70 | 1.013 | 8.82 | 11.91 | 1.013 | 10.11 | 18.32 | 1.013 | 5.41 | 16.36 | 1.013 | 3.00 | 17.28 |
| 1.019 | 9.68 | 3.54 | 1.019 | 13.66 | 17.67 | 1.019 | 16.06 | 27.79 | 1.019 | 8.16 | 24.28 | 1.019 | 3.94 | 24.33 |
| 1.025 | 12.93 | 4.72 | 1.025 | 19.45 | 23.20 | 1.025 | 24.63 | 38.59 | 1.025 | 11.59 | 33.14 | 1.025 | 5.11 | 31.47 |
| 1.031 | 17.01 | 6.14 | 1.031 | 27.21 | 30.27 | 1.031 | 36.91 | 53.24 | 1.031 | 16.10 | 43.14 | 1.031 | 6.77 | 39.30 |
| 1.038 | 22.62 | 7.97 | 1.038 | 38.38 | 39.25 | 1.038 | 55.25 | 71.20 | 1.038 | 21.61 | 54.36 | 1.038 | 8.79 | 47.74 |
| 1.044 | 31.55 | 10.21 | 1.044 | 53.89 | 50.07 | 1.044 | 80.88 | 94.46 | 1.044 | 28.99 | 67.80 | 1.044 | 11.37 | 57.02 |
| 1.050 | 44.58 | 13.31 | 1.050 | 74.58 | 62.77 | 1.050 | 114.03 | 122.29 | 1.050 | 38.82 | 84.24 | 1.050 | 14.41 | 67.33 |
| 1.056 | 62.93 | 17.64 | 1.056 | 103.79 | 79.64 | 1.056 | 157.77 | 156.84 | 1.056 | 52.57 | 103.39 | 1.056 | 18.01 | 78.80 |
| 1.063 | 86.50 | 23.21 | 1.063 | 140.76 | 99.45 | 1.063 | 213.88 | 198.11 | 1.063 | 69.17 | 126.07 | 1.063 | 23.21 | 91.84 |
| 1.069 | 116.72 | 29.97 | 1.069 | 187.30 | 124.63 | 1.069 | 283.90 | 247.19 | 1.069 | 90.47 | 153.03 | 1.069 | 29.22 | 107.24 |
| 1.075 | 155.29 | 39.31 | 1.075 | 247.71 | 156.25 | 1.075 | 370.58 | 306.65 | 1.075 | 119.61 | 185.94 | 1.075 | 36.77 | 124.38 |
| 1.081 | 204.50 | 51.40 | 1.081 | 322.51 | 193.98 | 1.081 | 473.17 | 374.49 | 1.081 | 155.46 | 225.04 | 1.081 | 45.75 | 145.31 |
| 1.088 | 269.71 | 67.25 | 1.088 | 421.10 | 240.98 | 1.088 | 598.90 | 456.81 | 1.088 | 203.65 | 275.28 | 1.088 | 57.79 | 171.06 |
| 1.094 | 357.00 | 87.38 | 1.094 | 548.29 | 301.55 | 1.094 | 753.88 | 561.17 | 1.094 | 270.62 | 340.53 | 1.094 | 74.18 | 204.12 |
| 1.100 | 484.04 | 115.51 | 1.100 | 724.81 | 385.76 | 1.100 | 966.14 | 714.12 | 1.100 | 365.55 | 437.14 | 1.100 | 96.21 | 256.15 |
| -45/+45 orientation<br>strip-x |  |  | -45/+45 orientation<br>off-x |  |  | -45/+45 orientation<br>equi-biax |  |  | -45/+45 orientation<br>off-y |  |  | -45/+45 orientation<br>strip-y |  |  |
| $\lambda_x : \lambda_y = 1.10 : 1.00$ | | | $\lambda_x : \lambda_y = 1.10 : 1.05$ | | | $\lambda_x : \lambda_y = 1.10 : 1.10$ | | | $\lambda_x : \lambda_y = 1.05 : 1.10$ | | | $\lambda_x : \lambda_y = 1.00 : 1.05$ | | |
| $\lambda_x$<br>[-] | $\sigma_x$<br>[kPa] | $\sigma_y$<br>[kPa] | $\lambda_x$<br>[-] | $\sigma_x$<br>[kPa] | $\sigma_y$<br>[kPa] | $\lambda_x$<br>[-] | $\sigma_x$<br>[kPa] | $\sigma_y$<br>[kPa] | $\lambda_y$<br>[-] | $\sigma_x$<br>[kPa] | $\sigma_y$<br>[kPa] | $\lambda_y$<br>[-] | $\sigma_x$<br>[kPa] | $\sigma_y$<br>[kPa] |
| 1.000 | 0.00 | 0.00 | 1.000 | 0.00 | 0.00 | 1.000 | 0.00 | 0.00 | 1.000 | 0.00 | 0.00 | 1.000 | 0.00 | 0.00 |
| 1.006 | 9.70 | 2.08 | 1.006 | 8.78 | 6.80 | 1.006 | 9.96 | 9.96 | 1.006 | 6.80 | 8.78 | 1.006 | 2.08 | 9.70 |
| 1.013 | 15.58 | 3.36 | 1.013 | 14.45 | 10.71 | 1.013 | 15.61 | 15.61 | 1.013 | 10.71 | 14.45 | 1.013 | 3.36 | 15.58 |
| 1.019 | 20.89 | 4.33 | 1.019 | 19.82 | 14.15 | 1.019 | 21.62 | 21.62 | 1.019 | 14.15 | 19.82 | 1.019 | 4.33 | 20.89 |
| 1.025 | 26.47 | 5.26 | 1.025 | 25.66 | 17.83 | 1.025 | 28.56 | 28.56 | 1.025 | 17.83 | 25.66 | 1.025 | 5.26 | 26.47 |
| 1.031 | 32.24 | 6.31 | 1.031 | 31.86 | 21.74 | 1.031 | 36.47 | 36.47 | 1.031 | 21.74 | 31.86 | 1.031 | 6.31 | 32.24 |
| 1.038 | 38.30 | 7.32 | 1.038 | 38.95 | 25.78 | 1.038 | 45.83 | 45.83 | 1.038 | 25.78 | 38.95 | 1.038 | 7.32 | 38.30 |
| 1.044 | 45.41 | 8.44 | 1.044 | 47.91 | 30.97 | 1.044 | 58.50 | 58.50 | 1.044 | 30.97 | 47.91 | 1.044 | 8.44 | 45.41 |
| 1.050 | 53.15 | 9.83 | 1.050 | 58.33 | 37.14 | 1.050 | 73.30 | 73.30 | 1.050 | 37.14 | 58.33 | 1.050 | 9.83 | 53.15 |
| 1.056 | 62.09 | 11.48 | 1.056 | 70.30 | 44.19 | 1.056 | 91.43 | 91.43 | 1.056 | 44.19 | 70.30 | 1.056 | 11.48 | 62.09 |
| 1.063 | 72.81 | 13.81 | 1.063 | 84.18 | 52.68 | 1.063 | 113.37 | 113.37 | 1.063 | 52.68 | 84.18 | 1.063 | 13.81 | 72.81 |
| 1.069 | 85.34 | 16.47 | 1.069 | 101.18 | 62.96 | 1.069 | 138.85 | 138.85 | 1.069 | 62.96 | 101.18 | 1.069 | 16.47 | 85.34 |
| 1.075 | 100.14 | 19.79 | 1.075 | 121.49 | 75.30 | 1.075 | 170.42 | 170.42 | 1.075 | 75.30 | 121.49 | 1.075 | 19.79 | 100.14 |
| 1.081 | 117.90 | 23.78 | 1.081 | 146.91 | 90.74 | 1.081 | 211.22 | 211.22 | 1.081 | 90.74 | 146.91 | 1.081 | 23.78 | 117.90 |
| 1.088 | 140.27 | 29.07 | 1.088 | 178.91 | 109.49 | 1.088 | 262.98 | 262.98 | 1.088 | 109.49 | 178.91 | 1.088 | 29.07 | 140.27 |
| 1.094 | 169.38 | 36.57 | 1.094 | 223.03 | 136.23 | 1.094 | 328.39 | 328.39 | 1.094 | 136.23 | 223.03 | 1.094 | 36.57 | 169.38 |
| 1.100 | 218.89 | 50.19 | 1.100 | 297.23 | 181.10 | 1.100 | 441.78 | 441.78 | 1.100 | 181.10 | 297.23 | 1.100 | 50.19 | 218.89 |

**Figure 5:**
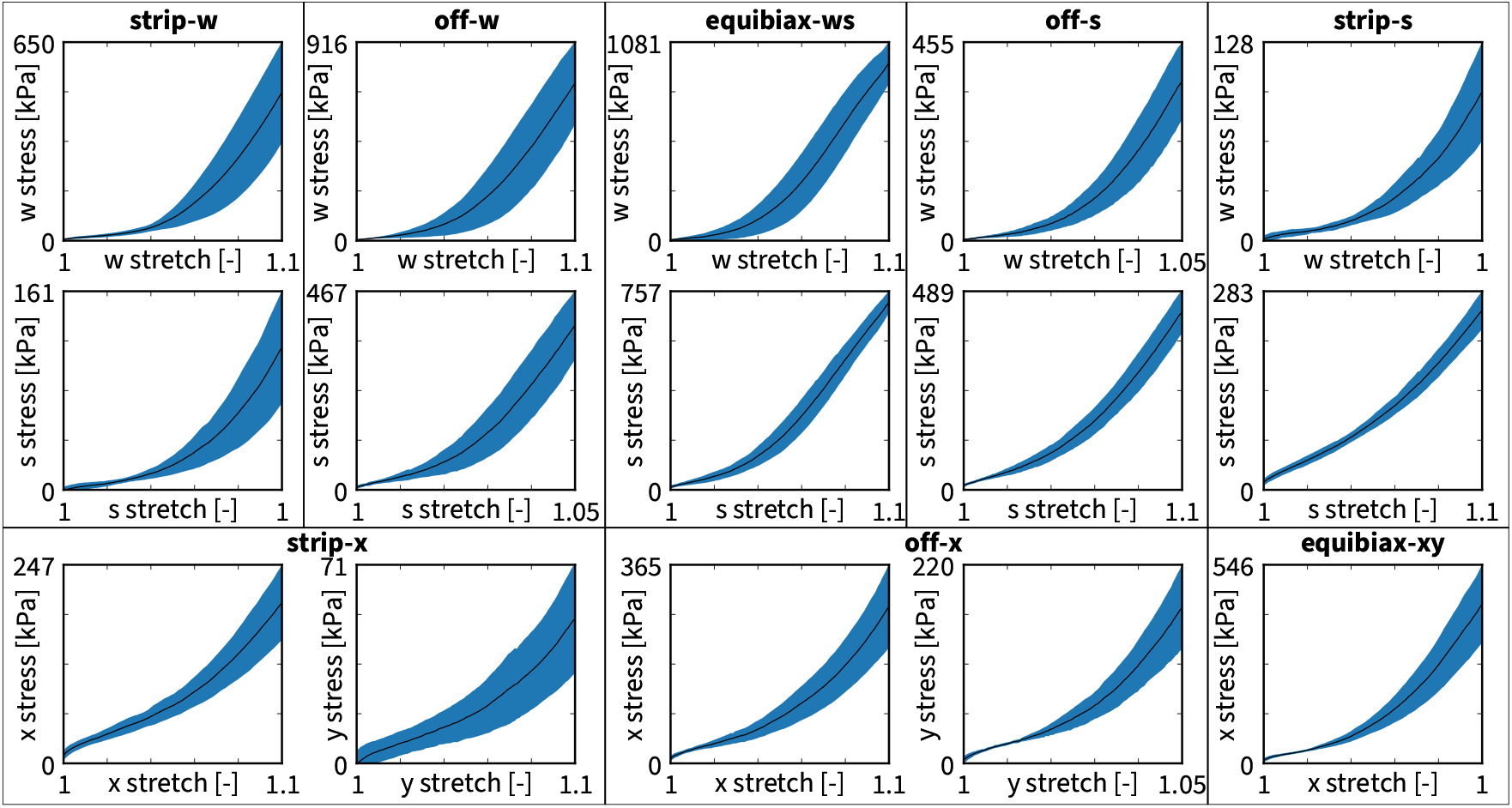
Biaxial test data during loading. Warp and shute stress-stretch data for strip-x, off-x, equibiax, off-y, and strip-y tests in the 0/90 orientation, top rows, and in the -45/+45 orientation, bottom row. Solid lines represent the means of *n* = 5 tests, shaded areas represent the standard deviations.

**Figure 6:**
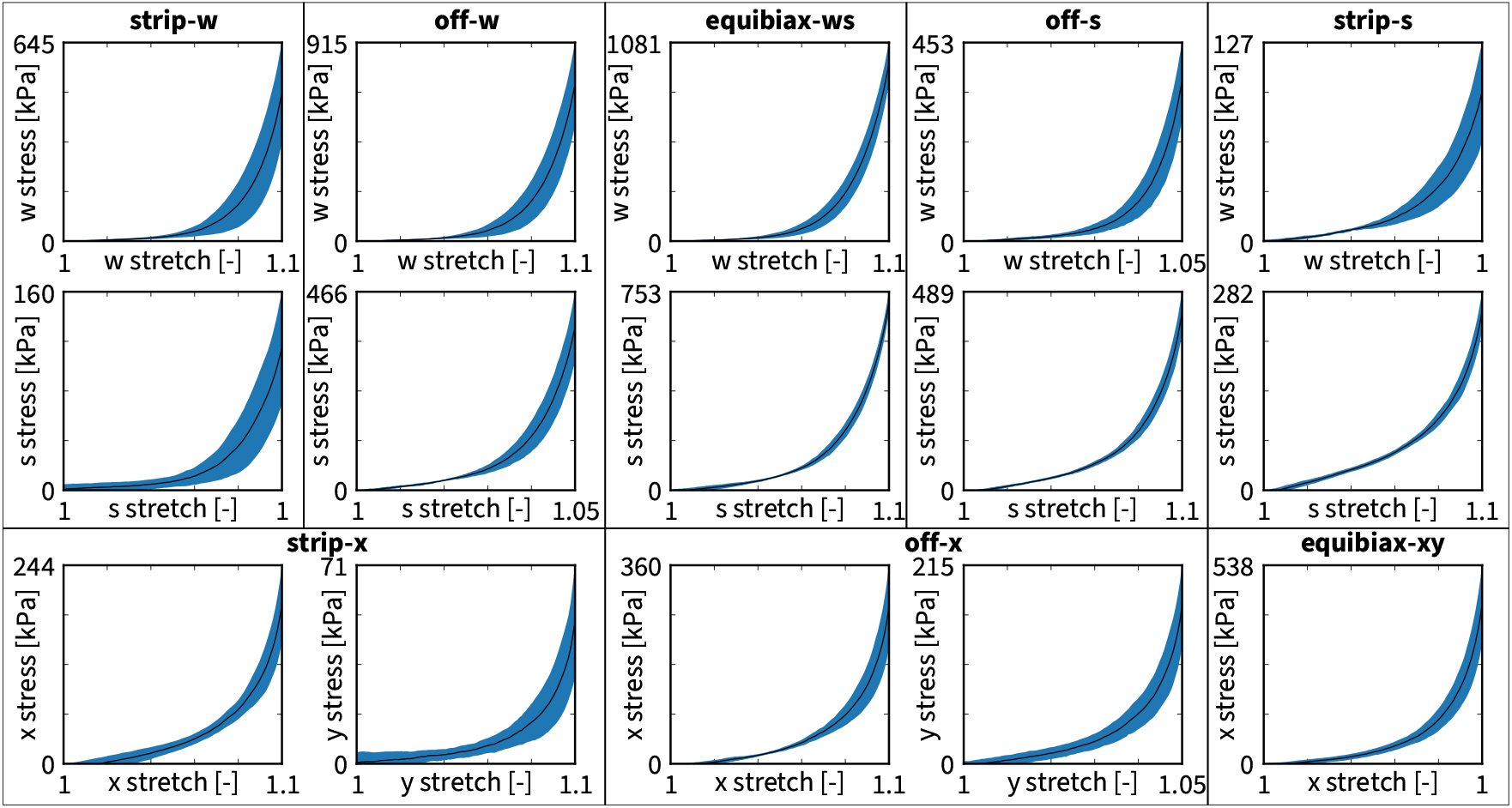
Biaxial test data during unloading. Stress-stretch data for strip-x, off-x, equibiax, off-y, and strip-y tests in the 0/90 orientation, top rows, and in the -45/+45 orientation, bottom row. Solid lines represent the means of *n* = 5 tests, shaded areas represent the standard deviations.

### 3.1. Mechanical signature of warp knitted fabric

All fifteen stress-stretch curves in Figures 5 and 6 display a similar trend: During the first half of the loading interval, the mesh behaves very compliant and the recorded stresses remain low; during the second half, the mesh stiffens and the stresses increase exponentially. Although present during both loading and unloading, this trend is less visible in the loading curves in Figure 5 than in the unloading curves in Figure 6, which also display smaller standard deviations. Intuitively, we expect the mesh to be stiffer in the warp than in the shute direction and Figures 5 and 6 and Table 2 confirm our expectation: In the strip-w and strip-s tests, the peak warp stress of 484 kPa is about twice as large as the peak shute stress of 256 kPa. In the equibiaxial test, the peak warp stress of 966 kPa is about one third larger than the peak shute stress of 714 kPa. We also expect the mesh to be stiffer in the 0/90 orientation, for which the loading axes are aligned with the warp and shute directions, than in the - 45/+45 orientation, for which the loading axes are rotated by 45 degrees against the warp and shute directions. In the strip-x and strip-y tests, the peak stress of 219 kPa is significantly lower than either of the two peak stresses of 484 kPa and 256 kPa of the unrotated setting. In the equibiaxial test, the peak stress of 442 kPa is also lower than two biaxial peak stresses of 966 kPa and 714 kPa of the unrotated setting. Taken together, our measurements confirm that knitted meshes display unique mechanical properties including an remarkable *initial flexibility*, a pronounced *nonlinear sti*ff*ening*, and an *extreme anisotropy*. To gain further insights into these characteristics, we now analyze the data using our constitutive neural networks and discover the two- and three-fiber models that best characterize our mesh.

### 3.2. Two-fiber architecture

First, we train the full network using the two-fiber architecture in Figure 3. We initialize the model parameters randomly and train the network without any regularization. Then, we use these trained parameters to re-initialize the parameters and train the network using *L*_0.5_ regularization with a regularization parameter *α* = 0.001. We use data from all experiments for training. Figure 7 shows the discovered model, with the contributions of each discovered stress term in a different color. The network discovers four non-zero terms, one is the exponential linear first invariant *I*_1_ Demiray term [8], one is a quadratic first invariant *I*_1_ term, and two are the exponential quadratic fourth warp and shute invariant *I*_4*w*_ and *I*_4*s*_ Holzapfel terms [16],

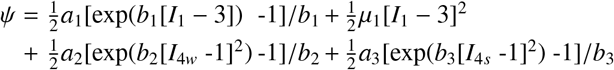

where the stiffness-like parameters are *a*_1_ = 91.24 kPa, µ_1_ = 3794 kPa, *a*_2_ = 3.31 kPa, *a*_3_ = 5.30 kPa, and the nonlinearity parameters are *b*_1_ = 0.17, *b*_2_ = 127.20, *b*_3_ = 94.68. Figure 7 illustrates the discovered two-fiber model along with the biaxial test data. It highlights four different terms in four characteristic colors and quantifies the goodness of all 15 fits in terms of the *R*^2^ values. The individual stress-stretch plots suggest that the discovered model performs fairly well on the experiments in the 0/90 orientation, but performs poorly on the strip-x experiment in the -45/+45 orientation, where it significantly underpredicts the x-stress and over-predicts the y-stress. This reduces the mean *R*^2^ value across all tests to 0.7573.

**Figure 7:**
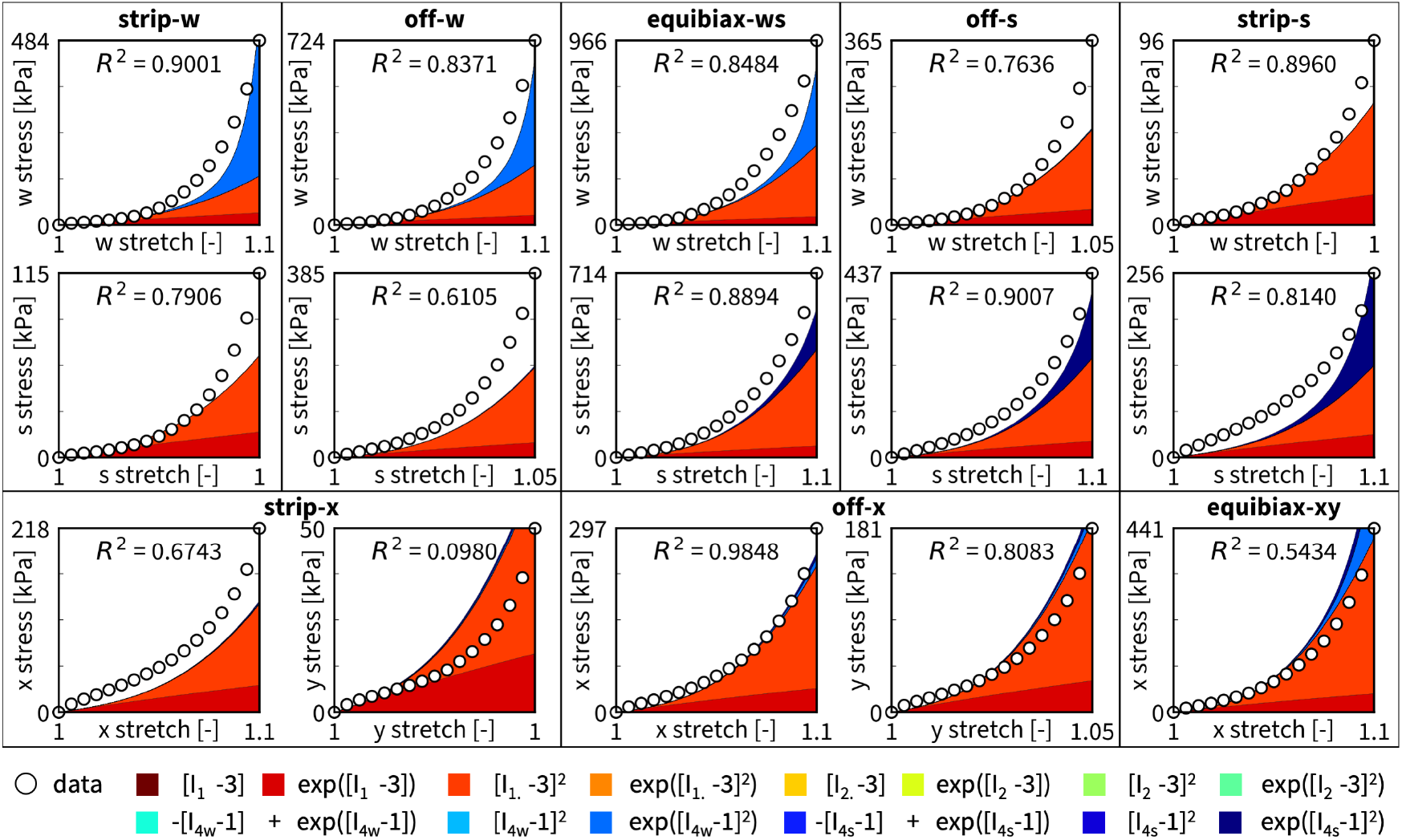
Discovered two-fiber model and biaxial test data. We train the two-fiber constitutive neural network from Figure 3 on all 15 data sets and sparsify the discovered model with *L*_0.5_ regularization. The network discovers a four-term model with two isotropic terms and two anisotropic terms, which we plot in their characteristic colors. The *R*^2^ values suggest that the discovered model performs decent on most of the tests in the 0/90 orientation, but performs particularly poorly on the strip-x experiment in the -45/+45 orientation where it underpredicts the x-stress and overpredicts the y-stress. The mean *R*^2^ value across all tests is 0.7573.

Additionally, we train a subset of the network in Figure 3 with *only anisotropic* terms. To do this, we constrain the weights of all eight isotropic terms to equal zero, *w*_1_, …, *w*_8_ = 0. We again use *L*_0.5_ regularization with a regularization parameter *α* = 0.001 and use all data for training. Figure 8 shows the discovered two-fiber model with no isotropic terms. The network discovers two anisotropic terms, both are the exponential linear Weiss terms [63], one in the fourth warp invariant *I*_4*w*_ and one in the fourth shute invariant *I*_4*s*_,

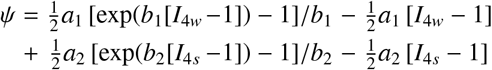

with the two stiffness-like parameters *a*_1_ = 2.998 kPa, *a*_2_ = 9.811 kPa, and the two nonlinearity parameters *b*_1_ = 21.07, *b*_2_ = 12.63. Notably, the fit of this model is extremely poor. The mean *R*^2^ value across all tests is as low as 0.5104, and five tests have *R*^2^ values of zero. We conclude that the orthotropic two-fiber model with no isotropic terms fails to accurately describe the behavior of the mesh.

**Figure 8:**
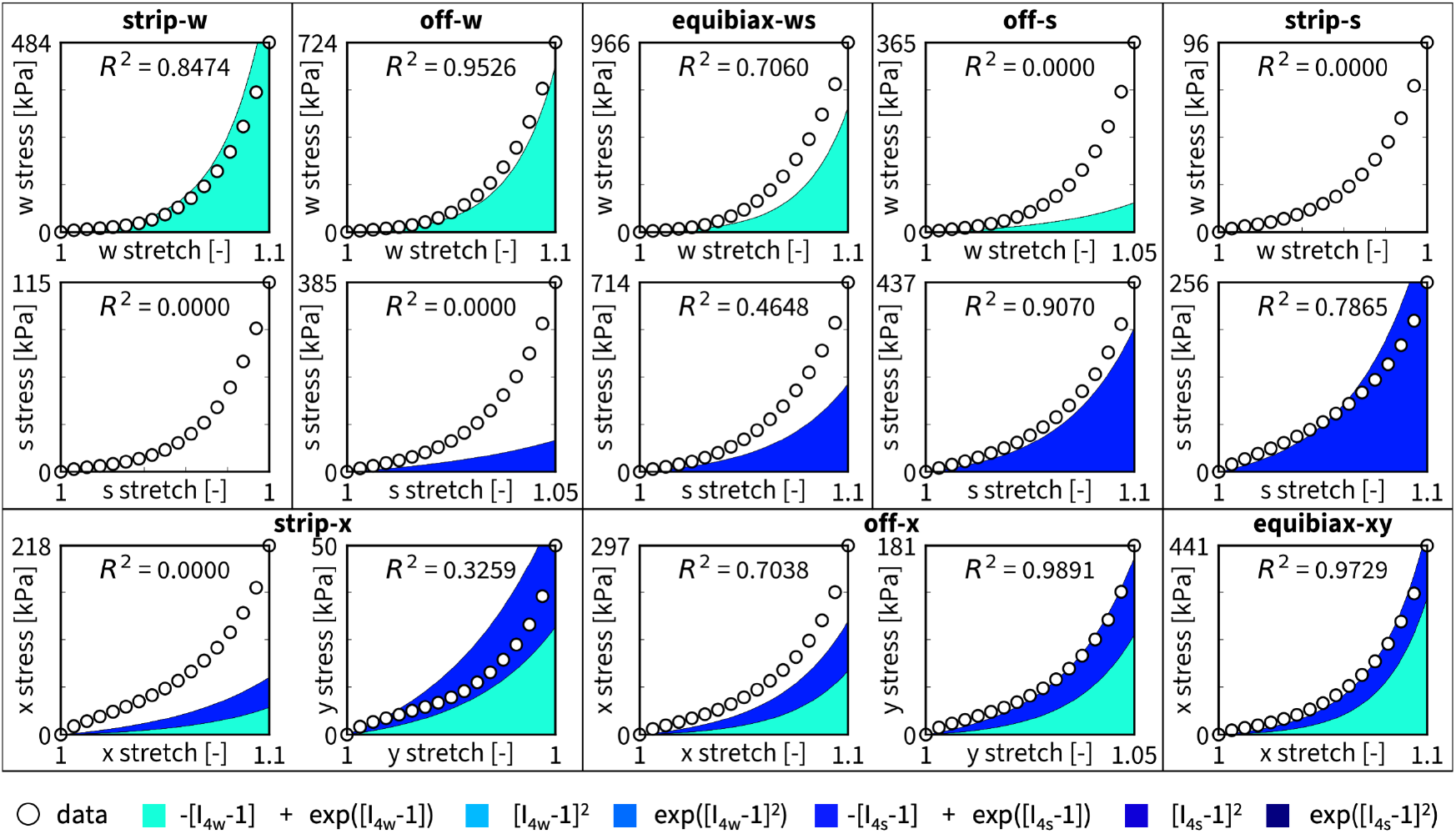
Discovered anisotropic two-fiber model and biaxial test data. We train the two-fiber constitutive neural network from Figure 4 on all 15 data sets and constrain the weights for the isotropic terms to be zero. The network discovers a two-term model with one term that is a function of the fourth warp invariant *I*_4*w*_ and one that is a function of the fourth shute invariant *I*_4*s*_, which we plot in their characteristic colors. The *R*^2^ values suggest that the discovered model performs poorly on most tests, with *R*^2^ values of zero for several tests. The mean *R*^2^ value across all tests is 0.5104.

Next, instead of sparsifying the model and reducing the number of terms with *L*_0.5_ regularization, we use *L*_0_ regularization to identify the best one- and two-term models by sampling all 14 single terms and all 91 pairs of two terms and minimizing the mean squared error of the 105 models. Figure 9 shows the minimized loss for all 105 combination of terms with the one-term models on the diagonale and the two-term models off the diagonale. For the two-fiber architecture, the best-in-class two-term model consists of one isotropic term, the exponential linear first invariant *I*_1_ Demiray [8], and one anisotropic term, the exponential quadratic fourth warp invariant *I*_4*w*_ Holzapfel term [16],

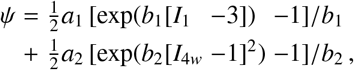

with the two stiffness-like parameters *a*_1_ = 25.20 kPa and *a*_2_ = 0.40 kPa, and the two nonlinearity parameters *b*_1_ = 17.30 and *b*_2_ = 127.30, and achieves a training loss of 0.077. Strikingly, while the data suggest that the mesh is highly anisotropic, the models with only anisotropic terms perform the worst.

**Figure 9:**
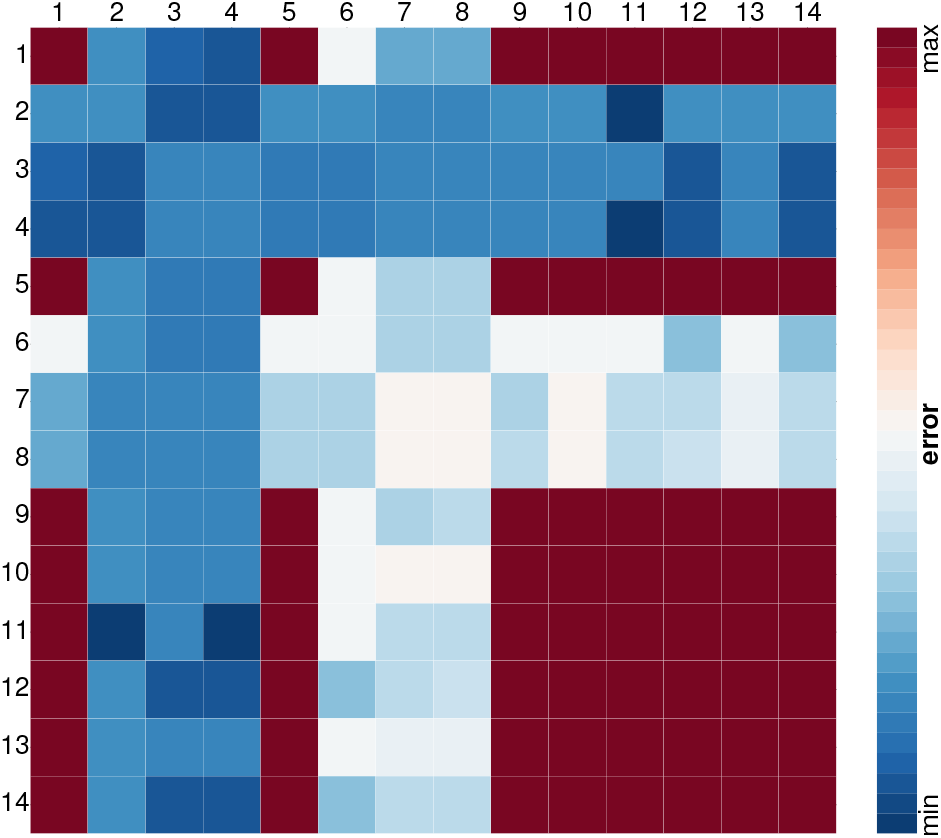
Best-in-class one- and two-term two-fiber models. All models are made up of 14 functional building blocks: linear, exponential linear, quadratic, and exponential quadratic terms of the first invariant *I*_1_, rows and columns 1 to 4, of the second invariant *I*_2_, rows and columns 5 to 8, of the fourth warp invariant *I*_4*w*_ rows and columns 9 to 11, and of the fourth shute invariant *I*_4*s*_ rows and columns 12 to 14. The color code indicates the mean squared error of the 14 one-term models on the diagonale, and of the 91 two-term models on the off-diagonale, ranging from dark blue, best fit, to dark red, worst fit.

### 3.3. Three-fiber architecture

Second, we train the full network using the three-fiber architecture in Figure 4. Again, we initialize the model parameters randomly and train the network without any regularization. Then, we use these trained parameters to re-initialize the parameters and train the network using *L*_0.5_ regularization with a regularization parameter *α* = 0.001. We use data from all experiments for training. Figure 10 shows the discovered model, with the contributions of each discovered stress term in a different color. The network discovers three non-zero terms, one is the exponential quadratic first invariant *I*_1_ Holzapfel term [16], and two are the exponential linear fourth warp and shute invariants *I*_4*w*_ and 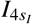 and 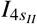 Weiss terms [63],

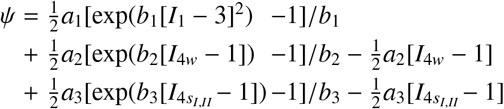

where the stiffness-like parameters are *a*_1_ = 2427 kPa, *a*_2_ = 0.16 kPa, *a*_3_ = 81.77 kPa, and the nonlinearity parameters are *b*_1_ = 0.51, *b*_2_ = 37.65, *b*_3_ = 4.78. Figure 10 shows the discovered three-fiber model along with the biaxial test data. Similar to the two-fiber model in Figure 7, the three-fiber model performs fairly well on the experiments in the 0/90 orientation. However, in contrast to the two-fiber model, the three-fiber model also performs well on the strip-x experiment in the +45/-45 orientation and achieves an *R*^2^ of 0.98 in the x-direction and 0.93 in the y-direction compared to the two-fiber model with only 0.67 in the x-direction and 0.10 in the y-direction. Its mean *R*^2^ value across all tests is 0.8614.

**Figure 10:**
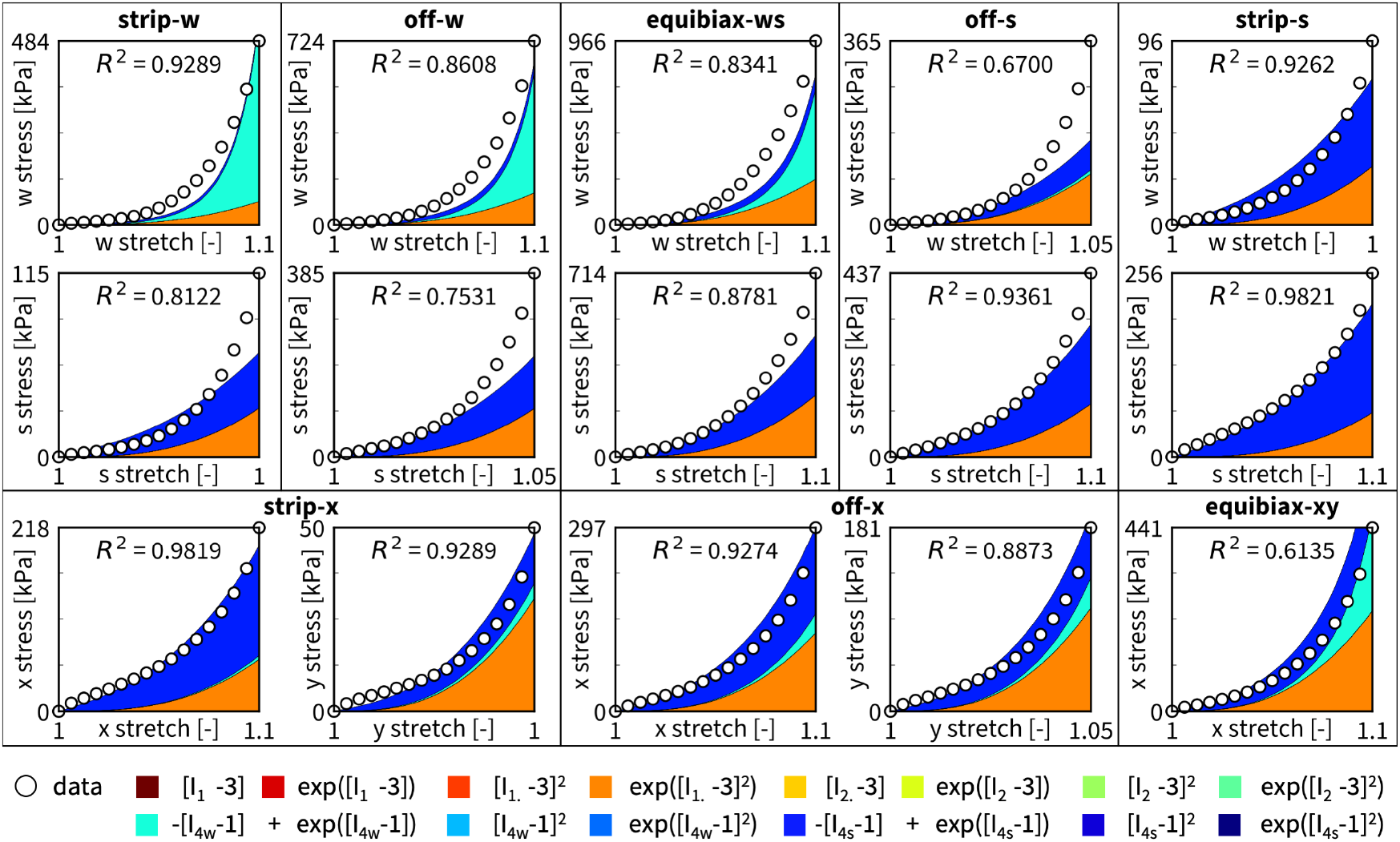
Discovered three-fiber model and biaxial test data. We train the three-fiber constitutive neural network from Figure 4 on all 15 data sets and sparsify the discovered model with *L*_0.5_ regularization. The network discovers a three-term model with one isotropic term and two anisotropic terms, which we plot in their characteristic colors. The *R*^2^ values suggest that the discovered model performs well on all fifteen tests. The mean *R*^2^ value across all tests is 0.8614.

Additionally, we train a subset of the network in Figure 4 with *only anisotropic* terms. Again, we constrain the weights of all eight isotropic terms to equal zero, *w*_1_, …, *w*_8_ = 0, use *L*_0.5_ regularization with *α* = 0.001, and use all data for training. Figure 11 shows the discovered three-fiber model with no isotropic terms. Similar to the two-term case, the network discovers two anisotropic terms, both are exponential linear Weiss terms [63], one in the fourth warp invariant *I*_4*w*_ and one in the fourth shute invariants 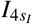 and 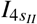,

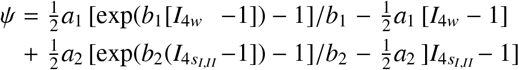

with the two stiffness-like parameters, *a*_1_ = 1.26 kPa, *a*_2_ = 28.93 kPa, and the two nonlinearity parameters *b*_1_ = 24.55, *b*_2_ = 7.20. Notably, the anisotropic three-fiber model in Figure 11 with a mean *R*^2^ value of 0.8199 provides a much better fit to the data than the anisotropic two-fiber model in Figure 8 with a mean *R*^2^ value of 0.5104, and, without any isotropic terms, performs almost as good as the three-fiber model in Figure 10 with a mean *R*^2^ value of 0.8614.

**Figure 11:**
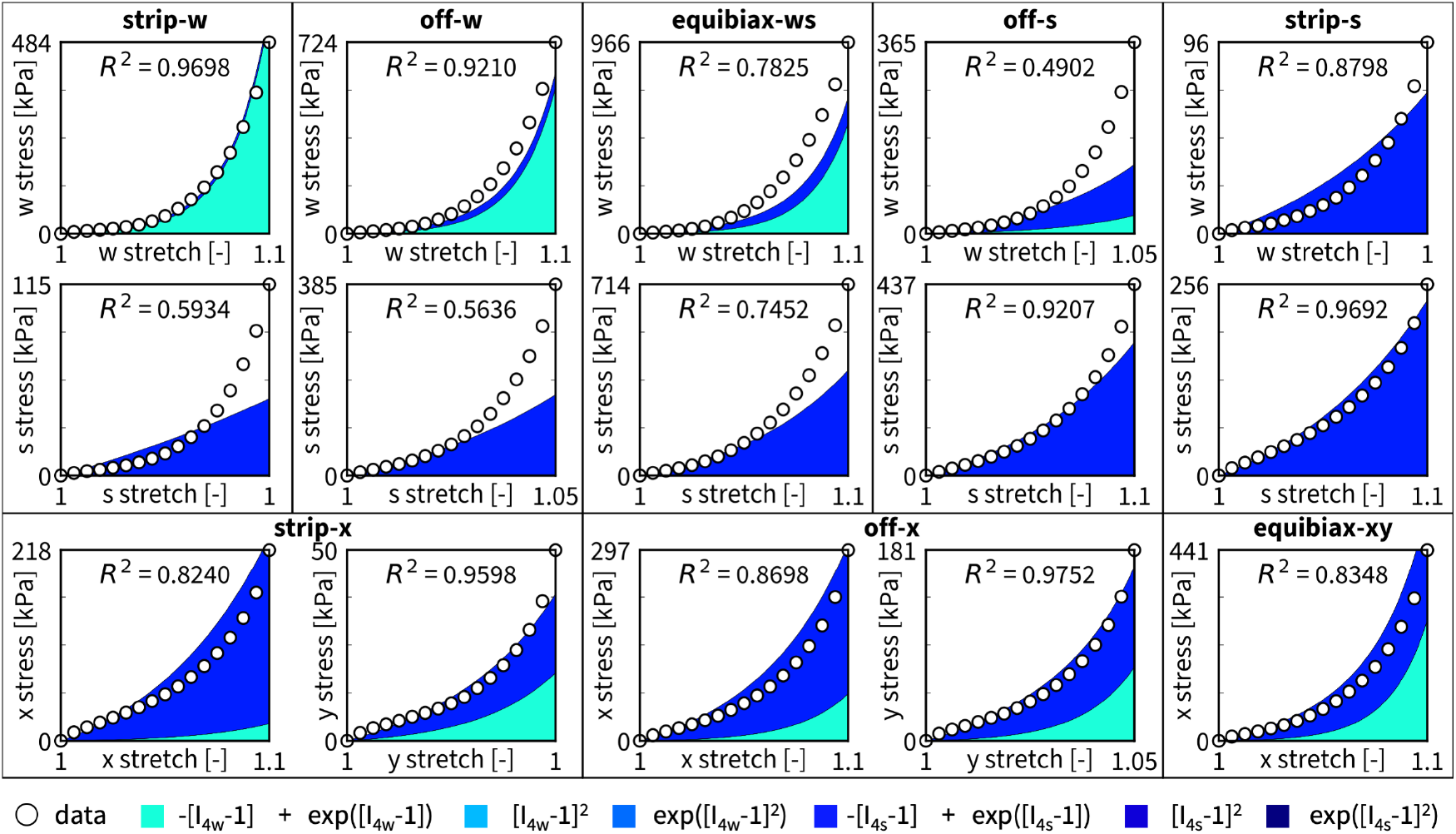
Discovered anisotropic three-fiber model and biaxial test data. We train the three-fiber constitutive neural network from Figure 4 on all 15 data sets and constrain the weights for the isotropic terms to be zero. The network discovers a two-term model with one term that is a function of the fourth warp invariant *I*_4*w*_ and one that is a function of the fourth shute invariants 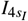 and 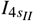, which we plot in their characteristic colors. The model that is discovered is very similar to the optimal two-term model shown in Figure 12. The *R*^2^ values suggest that, even without isotropic terms, the discovered model performs well across all fifteen tests. The mean *R*^2^ value across all tests is 0.8199.

Finally, we use *L*_0_ regularization to identify the best one- and two-term models by sampling all 14 single terms and all 91 pairs of two terms. Figure 12 shows the minimized loss for all 105 combination of terms. The first eleven terms in the three-fiber architecture are identical to the first eleven terms in the two-fiber architecture, which means the only difference between Figures 9 and 12 are the last three rows and columns associated with the shute invariants 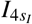 and 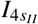. Interestingly, for the three-fiber architecture, the best-in-class two-term model is identical to the discovered non-isotropic model with two exponential linear Weiss terms [63], one in the fourth warp invariant *I*_4*w*_ and one in the fourth shute invariants 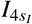 and 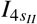,

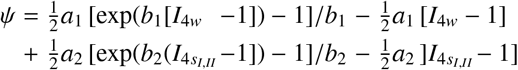

with the two stiffness-like parameters, *a*_1_ = 1.26 kPa, *a*_2_ = 28.93 kPa, and the two nonlinearity parameters *b*_1_ = 24.55, *b*_2_ = 7.20, and a training loss of 0.04. Strikingly, of all possible one- and two-term models, the best model characterizes the warp knitted mesh without any isotropic terms, and is exclusively made up of exponential linear terms of the squared stretches along the microstructural directions of the mesh.

**Figure 12:**
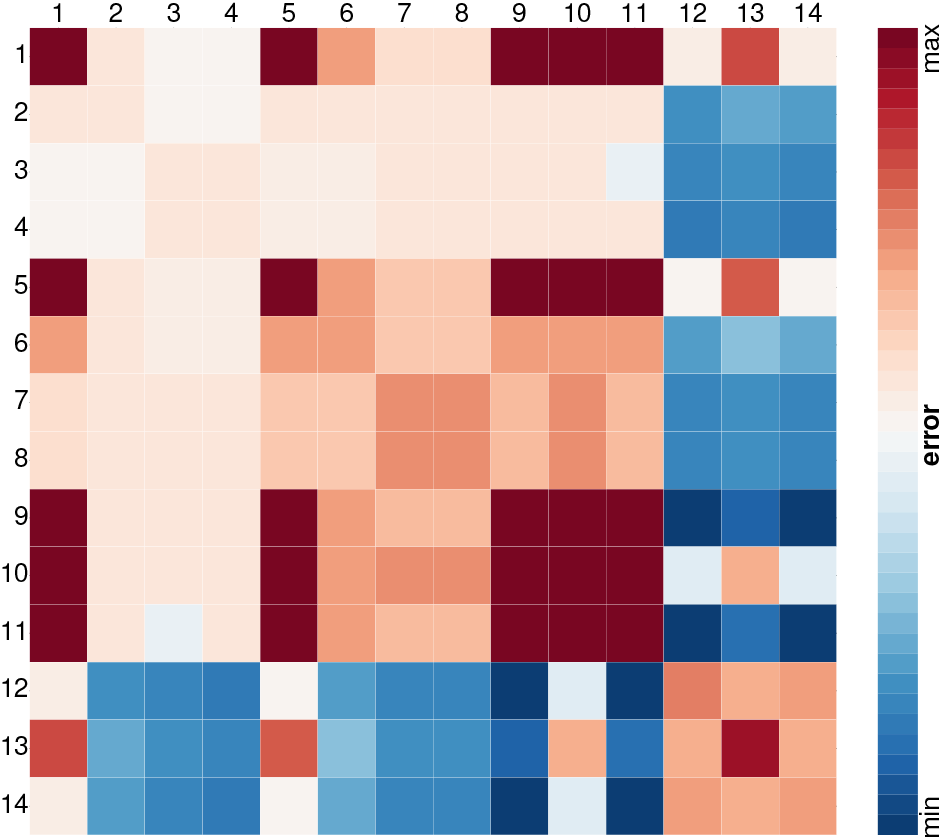
Best-in-class one- and two-term three-fiber models. All models are made up of 14 functional building blocks: linear, exponential linear, quadratic, and exponential quadratic terms of the first invariant *I*_1_, rows and columns 1 to 4, of the second invariant *I*_2_, rows and columns 5 to 8, of the fourth warp invariant *I*_4*w*_ rows and columns 9 to 11, and of the fourth shute invariants 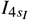 and 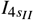 rows and columns 12 to 14. The color code indicates the mean squared error of the 14 one-term models on the diagonale, and of the 91 two-term models on the off-diagonale, ranging from dark blue, best fit, to dark red, worst fit.

### 3.4. Training and testing

To demonstrate that our discovered models have predictive capability beyond the data used in training, we train the network using a subset of the data and use the remaining data to test model performance. Yet, rather than randomly partitioning the available data into a training and test sets, we choose each of the tests to either be entirely part of the training set or test set. This is because the observations within a single test are highly correlated, and a constitutive model is most useful if it is capable of accurately predicting the resulting stress when the applied deformation takes a form that differs from the training data. We train the network with the three-fiber architecture from Figure 4 using four different training sets: the first training set consists of all experiments, the second consists of only experiments in the 0/90 orientation; the third consists of only experiments in the +45/-45 orientation; and the fourth consists of all experiments except strip-shute loading in the 0/90 orientation. Figure 13 shows the resulting coefficient of determination *R*^2^ for each stress-stretch curve when training the model on each of these four training sets. The figure illustrates that it is insufficient to train exclusively on the tests in the 0/90 orientation or on tests in the +45/-45 orientation; data from both orientations are essential to fully understand the constitutive behavior of our mesh. Notably, the model performance on the strip-shute tests when trained with all data except strip-shute is remarkably similar to the model performance when trained on all data. This suggests that the model is capable of extrapolating to unseen data, and shows that it does not overfit the training data.

**Figure 13:**
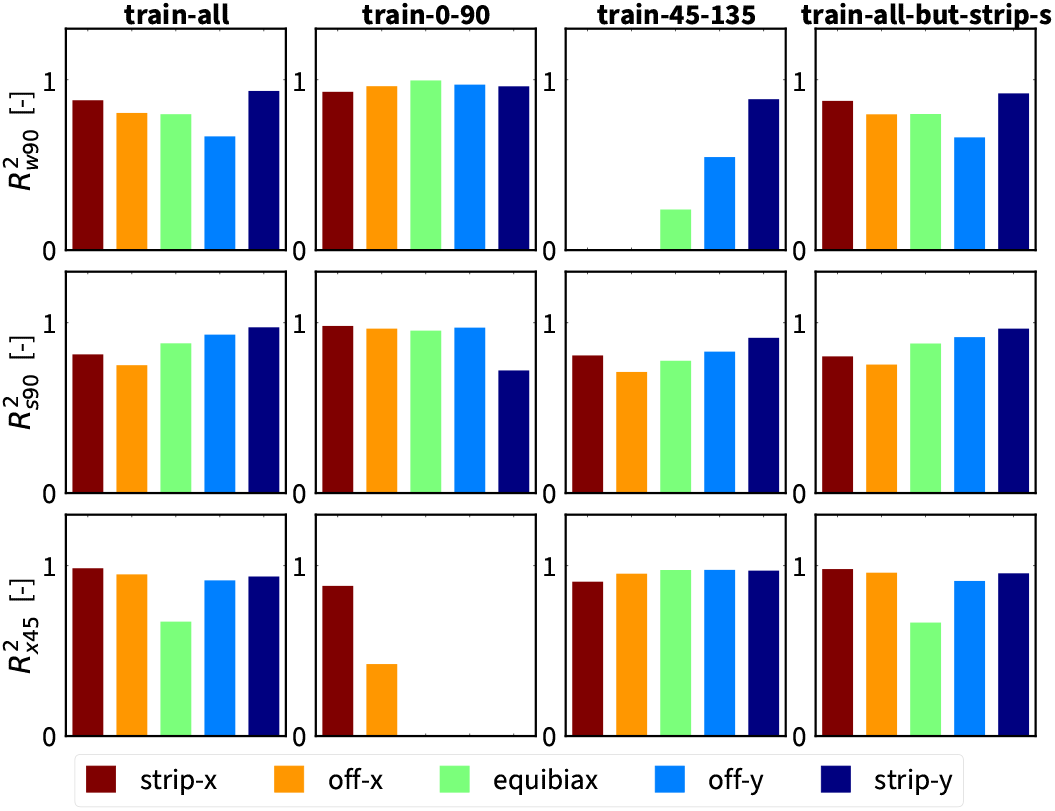
Performance of three-fiber network for four different training sets. The columns represent the four training sets: data from only the 0/90 orientation, data from only the +45/-45 orientation, all data, and all data except the strip-s data in the 0/90 orientation. The rows represent the coefficients of determination of the warp stress in the 0/90 orientation, the shute stress in the 0/90 orientation, and the x stress in the +45/-45 orientation. To accurately predict the stress in all loading conditions, the training set must include data from both the 0/90 and +45/-45 orientations.

## 4. Discussion

Synthetic meshes have unique mechanical properties, which make them ideal candidates for many engineering and medical applications. *The objective of this study was to provide insights into the mechanical signature of synthetic meshes using an integrative approach that combines biaxial testing and automated model discovery*. We prototyped this approach using a 0.5 mm thick warp knitted surgical mesh of extruded polypropylene. We tested the mesh in two different orientations, the 0/90 orientation with the loading axis aligned with the warp direction, and the -45/+45 orientation with the loading axis inclined by 45 degrees to the warp direction. We compared two families of microstructural models, a two-fiber model with a warp direction ***w*** and one orthogonal shute direction ***s***, and a three-fiber model with a warp direction ***w*** and two symmetrical shute directions ***s***_*I*_ and ***s***_*II*_ inclined by 60 degrees. Our study confirms our intuition that knitted meshes display a remarkable *initial flexibility*, a pronounced *nonlinear sti*ff*ening*, and an *extreme anisotropy*. Beyond these expected observations, it also reveals several exciting features of polypropylene meshes.

### Exponential linear fourth invariant terms dominate the constitutive response of polypropylene meshes

Throughout this study, we pursued different approaches to discover the best model and parameters to characterize textile structures. We trained a full neural network with fourteen independent terms and a subset of the network with only the six anisotropic terms, both for the two- and three-fiber model, with pronounced directions inclined by either 90 or 60 degrees. This allows us to discover the best model and parameters out of 2^14^ = 16, 384 and 2^6^ = 64 possible combinations of terms.

Strikingly, one term reoccured consistently through automated model discovery: the exponential linear fourth invariant term, 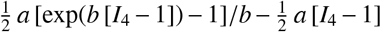. Originally proposed for soft biological tissues [63], this term is ideally suited to characterize the remarkable *initial flexibility* of the virgin mesh and the pronounced *nonlinear sti*ff*ening* as the loops of the mesh tighten upon loading. Of all four discovered models in Figures 7, 8, 10, and 11, three prominently feature this exponential linear fourth invariant term, both in the warp and shute directions *I*_4*w*_ and *I*_4*s*_, highlighted through the turquoise and blue colors.

We confirm the dominance of these two terms with an entirely independent best-in-class modeling study in Figures 9 and 12 that also identifies the the turquoise and blue terms to make up the best-in-class two-term model out of 91 possible two-term models, both for the two- and three-fiber microstructure. Interestingly, the full network model in Figure 10 with a mean *R*^2^ value of 0.8614 performs only marginally better than the reduced network model in Figure 11 with a mean *R*^2^ value of 0.8199. This suggest that we can confidently use the discovered two-term fourth-invariant model that features no isotropic terms to characterize the *ultra-anisotropic* nature of polypropylene meshes.

### Mixed invariant terms are critical to characterize the interaction of the warp and shute directions

To systematically explore the importance of the mixed invariant *I*_8*ws*_, we studied two different microstructural models. Figure 2 illustrates their two distinct microstructures, which represent the loops of the mesh as through warp direction ***w*** and the underlap through a single orthogonal shute direction ***s*** or two shoot directions ***s***_*I*_ and ***s***_*II*_, symmetrically offset by 60 degrees [21]. A direct comparison of the performance of our discovered two-fiber and three-fiber models in Figures 7 and 10 suggests that incorporating detailed microstructural information along all three directions is critical for model accuracy [57]. In Figure 7, we can see that the two-fiber model significantly underpredicts the x-stress and over-predicts the y-stress for the strip-x experiment in the +45/-45 orientation. To show that this error is a direct consequence of the two-fiber model, we take a closer look at the anisotropic stress contributions. In particular, when the strain energy is not a function of *I*_8*ws*_, the contribution of the anisotropic terms to the x- and y-stress in the +45/-45 orientation is 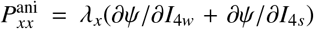 and 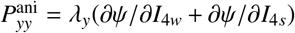, thus, 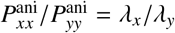 At maximum strip-x loading in the +45/-45 orientation, the stretches are *λ*_*x*_ = 1.1 and *λ*_*y*_ = 1.0, such that *λ*_*x*_/*λ*_*y*_ = 1.1, and the stresses are 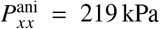 and 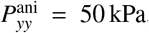, such that 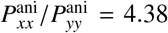. We can directly see this discrepancy in Figure 8, where the discovered model has an *R*^2^ value of zero for the x-stress in strip-x loading in the +45/-45 orientation. This suggests that an appropriate model for warp knitted meshes should indeed be a function of the mixed invariant *I*_8*ws*_. If the strain energy *ψ* is a function of the mixed invariant *I*_8*ws*_ [34], the derivative *∂ψ*/*∂I*_8*ws*_ contributes a positive component to 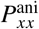 and a negative component to 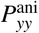. For positive mixed invariants, *∂ψ*/*∂I*_8*ws*_ > 0, such that 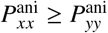, and thus 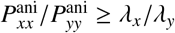, which is what we observe in Table 2. This observation is in line with several previous studies that have acknowledged the importance of the mixed invariant for double-fiber reinforced nonlinear elastic materials [36].

### Understanding the shear response is critical to model textile structures

Even with an appropriate constitutive neural network that is informed by the microstructure of the mesh, our inability to measure shear strain in biaxial loading [10] results in a loss of model accuracy in certain loading modes. Most obviously, when comparing Figures 7 and 10, we notice that the model generally under-predicts the shute and warp stresses in the 0/90 orientation, and over-predicts the x- and y-stresses in the +45/-45 orientation. When taking a closer look at the states of maximum deformation in equibiaxial loading, we see that when *λ*_*w*_ = *λ*_*s*_ = *λ* = 1.1 in the 0/90 orientation, the invariants and mixed invariants are identical to when *λ*_*x*_ = *λ*_*y*_ = *λ* = 1.1 in the +45/-45 orientation. In both cases, *I*_1_ = 2*λ*^2^ + *λ*^−4^ and *I*_2_ = 2*λ*^−2^ + *λ*^4^ and *I*_4*w*_ = *I*_4*s*_ = *λ*^2^ and *I*_8*ws*_ = 0. As a result, the strain energy *ψ* and its partial derivatives are identical in both cases. Thus, 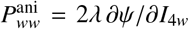 and 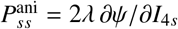 and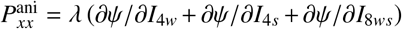.

Since *I*_8*ws*_ = 0 and *ψ* is an even function of *I*_8*ws*_, by symmetry, we know that *∂ψ*/*∂I*_8*ws*_ = 0, and thus, *P*_*ww*_ + *P*_*ss*_ = 2 *P*_*xx*_. However, looking at the data, *P*_*ww*_ + *P*_*ss*_ = 996 kPa + 714 kPa = 1710 kPa, while 2 *P*_*xx*_ = 884 kPa. This contradiction is triggered by our assumptions in equations (5) and (16) that the deformation remains *homogeneous* and *shear free* at all times, such that the deformation gradient ***F*** remains diagonal, 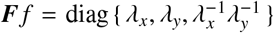. This condition holds for samples loaded in biaxial tension, if their microstructure is symmetric about the two loading directions. While this is true for isotropic materials in general, and it holds for orthotropic materials in the 0/90 configuration, it does not always hold for orthotropic materials in the +45/-45 configuration: Loading non-symmetrically mounted orthotropic materials in biaxial tension tests may actually result in non-zero shear strains [52]. Since the CellScale BioTester 5000 grips the material using slender tines that are stiff in tension but compliant in bending, a more accurate boundary condition would be that the axial stretches are equal to the values measured by the device, and the shear stresses are equal to zero. In this case, the shear strains would be non-zero and would take the values that minimize the strain energy given the prescribed axial stretches in the two loading directions. This change would decrease the predicted strain energy in the +45/-45 orientation, while leaving the strain energy in the 0/90 orientation unchanged. As a result, we would expect 2*P*_*xx*_ < *P*_*ww*_ + *P*_*ss*_, which is indeed what we observe in Table 2. To accurately incorporate this effect, we would need to measure the shear strain, which the CellScale BioTester 5000 does not directly control or measure.

### Our networks reliably extrapolate to unseen data

Our study confirms, that with an appropriate microstructural representation, our constitutive neural network discovers generalizable anisotropic constitutive models, provided that we use training data from both the 0/90 and +45/-45 orientations. From the goodness-of-fit bar plots in Figure 13, it is clear that training on only data from the 0/90 orientation results in inaccurate model predictions in the +45/-45 orientation, and vice versa. This result is not surprising since certain characteristics of the material are only active and visible in one of the two orientations. In particular, in the +45/-45 orientation, by symmetry *I*_4*w*_ = *I*_4*s*_ for all loading states, which means that it is not possible to independently probe the *I*_4*w*_ and *I*_4*s*_ terms. Similarly, in the 0/90 orientation, *I*_8*ws*_ = 0 for all loading states of the two-fiber models, so we cannot measure the effect of *I*_8*ws*_ on the stress. Thus, the training data must consist of data from both orientations for the model to robustly predict stresses under all possible conditions. Another interesting observation from Figure 13 is that, when training on all the available data except the strip-shute data in the 0/90 configuration, the discovered model achieves similar *R*^2^ values for the training and test data when compared to the model trained on all the data. This suggests that our model is not overfitting the data, and that when trained on a subset of data from both orientations, it is able to accurately predict the stresses for deformation states that we do not use in training.

### From model discovery to generative material design

Our constitutive neural networks in Figures 3 and 4 solve the forward problem to *discover the best model and parameters* that describe a given material, in our case the textile microstructure. Inversely, we could use our constitutive neural networks and solve the inverse problem to *discover the best material* for a given model and parameters, for example, a desired directional stiffness. Both problems combined represent a classical example for bidirectional learning where layer-wise relevance propagation can provide insight into the forward problem of model discovery, whereas layer-wise relevance backpropagation can provide insight into the inverse problem of material discovery [2]. Alternatively, recent advances in structural feature representation and generative neural networks now allow us to more efficiently design materials with tailored properties and functions. In materials science, generative neural networks are currently gaining immense popularity in the design of solid-state crystalline materials [65], where the features that represent the crystalline microstructure are the atom type, the lattice vectors, and the atomic coordinates in the Euclidian space [43]. In textile science, these features naturally translate into yarn type, yarn angles, and knot or loop coordinates. Two popular and emerging models for inverse material design are variational autoencoders and generative adversarial networks [64]. A *variational autoencoder* consists of an encoder that transforms the input sample feature vector into the latent space where it generates the latent space vector *z* from a normal distribution *N*(σ, µ) and a decoder that reconstructs the sample from the given hidden distribution. A *generative adversarial network* consists of a generator that generates samples from random noise variables and a discriminator that determines whether a sample is valid or invalid. Figure 14 compares the material design process with our constitutive neural network for forward model discovery and inverse material discovery with these two popular generative neural networks, variational autoencoders and generative adversarial networks. Adapting neural network modeling to design programmable textile metamaterials with tunable properties and functions would open unique opportunities in textile science with possible applications to wearable devices, stretchable electronics, and smart fabrics.

**Figure 14:**
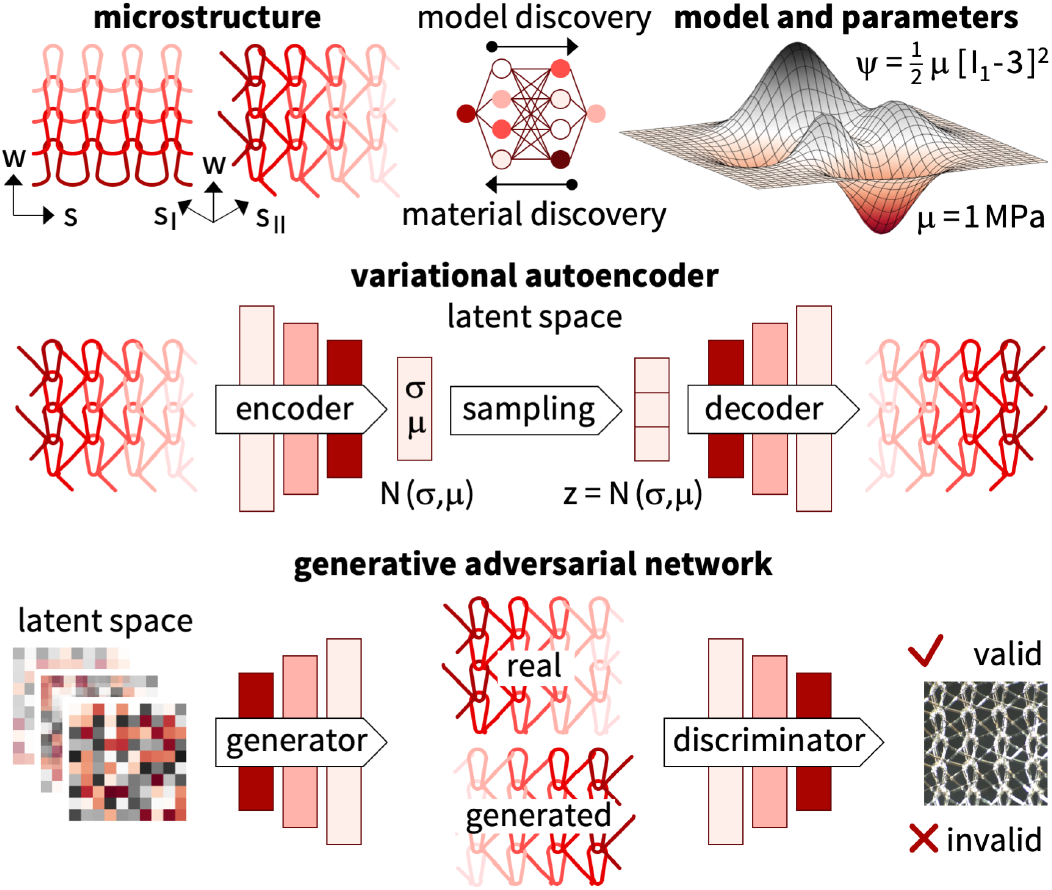
Material design with generative neural networks. Constitutive neural networks solve the forward problem to *discover models and parameters* for given microstructures, and could solve the inverse problem to *discover materials* for given models and parameters. A *variational autoencoder* for generative material design combines an *encoder* to transform the input sample feature vector into the latent space where it generates the latent space vector *z* from a normal distribution *N*(σ, µ) and a *decoder* to reconstruct the sample from the given hidden distribution. A *generative adversarial network* for generative material design combines a *generator* to generate samples from random noise variables and a *discriminator* to determine whether a sample is valid or invalid.

### Limitations and future directions

Our study presents a first step in characterizing textile structures using automated model discovery. Here we prototype this approach for a single warp knitted polypropylene mesh. While our experience with other material types [27, 28, 30, 48] suggests that our approach will generalize naturally to other textile structures–woven, werf knitted or warp knitted–our current study has a few limitations that point towards future research directions: First, our current study is limited to the hyperelastic regime. However, we have recorded separate loading and unloading data and expanding the model discovery process the inelastic regime using viscoelastic [62] or general inelastic [15] constitutive artificial neural networks would be the next logical step. Second, our study focuses on characterizing the two-dimensional in-plane behavior or the fabric structure. Expanding it to a more physiological fully three-dimensional in- and out-of-plane characterization is conceptually possible, but would require additional tests, for example, the ball burst test that characterizes the indentation response of the fabric as a thin membrane [6]. Third, our biaxial test setup uses square samples mounted by tines or rakes [10]. Alternatively, we could have mounted the samples using clamps, and we plan to investigate the impact of different mounting techniques in a future study. Finally, our current study assumes a homogeneous shear free state. A possible future extension would be to quantify shear strains using full field data from digital image correlation [1] and embed model discovery within the solution of real boundary value problems with possibly heterogeneous stresses and stretches [11].

## 5 Conclusion

Characterizing the mechanical properties of synthetic meshes is critical to understand their unique properties and functions. To date, identifying appropriate constitutive models for woven and knitted textiles poses a critical barrier to mechanically tailoring and fine-tuning these structures to individual needs. Machine-learning approaches can discover anisotropic constitutive models from biaxial data; yet, existing approaches are limited to training data from a single mounting orientation.

Here we show that this approach can result in superficial constitutive models that generalize poorly to unseen data. In contrast, the new approach we advocate here uses data from at least two different mounting orientations, and robustly discovers models that perform well during both training and testing. Importantly, our study shows that the accuracy of the discovered models is highly sensitive to an accurate representation of the microstructural architecture of the sample: Even if the textile fabric appears orthotropic at first glance, an accurate kinematic characterization of both warp and shute directions is critical to discover robust and reliable models. We demonstrate that these models are dominated by exponential linear fourth invariant terms that uniquely capture the remarkable initial flexibility, pronounced nonlinear stiffening, and extreme anisotropy of warp knitted polypropylene meshes. We anticipate that the tools we have developed here will generalize naturally to other textile fabrics–woven or knitted, weft knit or warp knit, with laid-in stitches or plain, polymeric or metallic–and, ultimately, will enable the robust discovery of anisotropic constitutive models for a wide variety of textile structures. Beyond discovering constitutive models, we envision to exploit automated model discovery as a novel strategy for the generative material design of wearable devices, stretchable electronics, and smart fabrics, as programmable textile metamaterials with tunable properties and functions.

## Data availability

Our source code, data, and examples are available at https://github.com/LivingMatterLab/CANN.

## Acknowledgments

This work was supported by the NSF Graduate Student Fellowship to Jeremy A. McCulloch and by the NSF CMMI Award 2320933 Automated Model Discovery for Soft Matter and by the ERC Advanced Grant 101141626 DISCOVER to Ellen Kuhl.

